# Antimicrobial peptides do not directly contribute to aging in *Drosophila*, but improve lifespan by preventing dysbiosis

**DOI:** 10.1101/2022.08.24.504952

**Authors:** M.A. Hanson, B. Lemaitre

## Abstract

Antimicrobial peptides (AMPs) are innate immune effectors first studied for their role in host defense against bacterial and fungal infections. Recent studies have implicated these peptides in the clearance of aberrant cells and various neurological processes including neurodegenerative syndromes. In *Drosophila*, an array of AMPs are produced downstream of the Toll and Imd NF-κB pathways in response to infection. Many studies have suggested a role for the Imd pathway and AMPs in aging in this insect, supported by the upregulation of AMPs with aging (so-called “inflammaging”). However, functional studies using RNAi or over-expression have been inconclusive on whether and how these immune effectors impact aging.

Leveraging a new set of single and compound AMP gene deletions in a controlled genetic background, we have investigated how AMPs contribute to aging. Overall, we found no major effect of individual AMPs on lifespan, with a possible exception of *Defensin*. However, *ΔAMP14* flies lacking 14 AMP genes from seven families display a reduced lifespan. Interestingly, we found an increased bacterial load in the food medium of aged *ΔAMP14* flies, suggesting that the lifespan reduction of these flies was due to a failure in controlling the microbiome. Consistent with this idea, use of germ-free conditions extends the lifespan of *ΔAMP14* flies. Overall, our results do not point to an overt role of individual AMPs in lifespan. Instead, we find that AMPs collectively impact lifespan by preventing dysbiosis over aging. This is consistent with our previous study showing that AMPs control the gut microbiome, and many works showing that dysbiosis is detrimental upon aging. In the course of our experiments, we also uncovered a strong impact of a *Drosophila nora virus* infection on lifespan, and share our experience in reconciling our data given this confounding cryptic factor.

## Introduction

Antimicrobial peptides (AMPs) are innate immune effectors found in plants and animals that show microbicidal activity in vitro. They are typically cationic and amphipathic, and disrupt microbial membranes that are more negatively charged (Broekaert et al., 1997; Christensen et al., 1988; Ludtke et al., 1996; Zasloff, 1987). Recent studies in various animal models have pointed to a role for AMPs beyond microbial infection, notably in aging and aging-related diseases (Deslouches and Di, 2017; Mookherjee et al., 2020; Semple and Dorin, 2012; Smith and Gwyer Findlay, 2022). Like other animals, AMP genes are upregulated upon aging in *Drosophila*, and disparate studies have suggested a role for AMPs as causative agents of aging, neurodegeneration or mitochondrial stress (Arora and Ligoxygakis, 2020; Garschall and Flatt, 2018; Hanson and Lemaitre, 2020). Here we have leveraged a set of isogenic AMP deficient flies to analyze the contribution of AMPs to aging in *Drosophila*.

Antimicrobial peptides are well characterized in *Drosophila* for their contribution to host defence. There are currently eight families of inducible AMPs known in *D. melanogaster*: the antifungals *Drosomycin, Baramicin*, and *Metchnikowin*; *Cecropins* (four inducible genes) and *Defensin* that have both antibacterial and some antifungal activities; and *Drosocin, Attacins* (four genes) and *Diptericins* (two genes) that primarily exhibit antibacterial activity (Hanson et al., 2021; Imler and Bulet, 2005). In addition, the *Drosophila* genome encodes many other host defence peptide families such as *Daisho* (two genes) and *Bomanin* (12 genes), where overt antimicrobial activity in vitro is not yet demonstrated, but functional studies have shown they are important in vivo to resist microbial infection (Clemmons et al., 2015; Cohen et al., 2020). While most *Drosophila* defence peptide genes are strongly induced in the fat body in response to systemic infections, many show specific and complex patterns of expression in tissues such as the trachea, gut, salivary glands, or reproductive tracts (Ferrandon et al., 1998; Tzou et al., 2000).

AMPs genes are regulated at the transcriptional level by the Toll and Imd NF-kB signaling pathways upon systemic infection, or by the Imd pathway in local epithelia (Lemaitre and Hoffmann, 2007; Myllymäki et al., 2014). It is well established that Imd and Toll deficient flies show marked susceptibility to microbial infection. Until recently, the importance of immune effectors downstream of these pathways, notably AMPs, was unclear. With the advent of CRISPR gene editing, we systematically deleted seven families of AMP genes (*Defensin, Cecropin, Drosocin, Attacin, Diptericin, Drosomycin*, and *Metchnikowin*) and analyzed their contributions to host defence individually or collectively. We found that *Drosophila* AMPs are essential downstream of the Imd pathway to resist systemic infection by Gram-negative bacteria. *Drosophila* AMPs also contribute downstream of the Toll pathway to combat fungi and to a lesser extent Gram-positive bacterial infection, although *Bomanins* play a more prominent role against these micro-organisms (Carboni et al., 2021; Clemmons et al., 2015; Hanson et al., 2019a). Use of fly lines carrying combinations of AMP mutations revealed that they can function either additively or synergistically against some microbes, but in some cases AMPs exhibit striking specificity, with one peptide contributing most of the AMP-dependent defence against a specific pathogen (Hanson et al., 2019a; Hanson et al., 2022a; Hanson et al., 2022b; Unckless et al., 2016). *Drosophila* AMPs are also important to control the fly microbiome downstream of the Imd pathway, particularly for their role in regulating Gram-negative bacteria like *Acetobacter* (Marra et al., 2021).

While *Drosophila* AMPs were initially investigated for their contribution to host defence, AMP upregulation is also observed in non-immune contexts, including anti-tumor defence (Araki et al., 2019; Krautz et al., 2020; Parvy et al., 2019), neurodegeneration (Cao et al., 2013; Kounatidis et al., 2017; Petersen et al., 2013; Shukla et al., 2019; van Alphen et al., 2022), and aging (Glittenberg and Silas, SukritGlittenberg, M. T., Silas, S., MacCallum, D. M., Gow, N. a R., & Ligoxygakis, 2011; Lai et al., 2007). This suggests that AMPs could have roles beyond their traditional function as microbicidal agents. Notably, transcriptomic studies revealed that high AMP expression, reflecting an increase in Imd pathway activity, is a hallmark of aging in flies (Landis et al., 2004; Pletcher et al., 2002; Seroude et al., 2002; Zerofsky et al., 2005). This is reminiscent of the situation observed in humans where a low-grade chronic inflammation is observed upon aging, termed “inflammaging” (Franceschi et al., 2000; Franceschi et al., 2017; Liang and Diana, 2020). The key question now is to decipher if this high and chronic activation of the immune response is simply a marker of aging, or if cytotoxic immune effectors like AMPs accelerate aging-associated syndromes.

To address this question, several reports have investigated the role of the Imd pathway and AMPs in aging in *Drosophila* with contrasting outcomes. Most of these studies targeting Imd itself, or the Imd transcription factor *Relish* report that Imd pathway mutants are short-lived, (Cai et al., 2021; Kounatidis et al., 2017; Petersen et al., 2013). However one study suggested that Imd mutation itself improves lifespan, as did fat body or whole-body AMP knockdown (Lin et al., 2018). The negative impact of Imd pathway downregulation has been associated with defects in gut homeostasis (Buchon et al., 2009), exaggerating the decline in gut resilience upon aging (Rera et al., 2012), leading to invasion of microbes into the hemolymph that drive mortality (Clark et al., 2015). Indeed the guts of aged *Relish* mutant flies display precocious loss of compartmentalization, increased permeability, and dysplasia (Liu et al., 2022). However preventing the over-activation of Imd in the gut through transgene expression can improve lifespan, suggesting a careful balance of immune signalling in the gut is needed for optimal health with aging (Cai et al., 2021; Guo et al., 2014; Iatsenko et al., 2018). Some of these symptoms could be reverted when flies were raised axenically pointing to a role of the microbiome in the precocious aging of the gut (Buchon et al., 2009; Clark et al., 2015; Iatsenko et al., 2018; Liu et al., 2022; Zhu et al., 2021). Another study reports that AMP genes are strongly induced in the head of old flies, and that silencing *Relish* in glia can extend life span (Kounatidis et al., 2017). This would suggest that Imd-mediated immune responses drive aging by directly affecting brain activity. Indeed, many studies report that suppression of the Imd pathway can prevent neurodegeneration in various neurodegenerative disease models. For instance, flies mutated in the serine/threonine kinase *ataxia telangiectasia mutated* (*ATM*, also called *tefu*) display reduced lifespan associated with increased neuronal lesions, which can be rescued by combination with *Relish* mutation (Petersen et al., 2013). *Relish* deletion also rescues the aging-dependent neurodegeneration of *Cdk5-alpha* mutant flies (Shukla et al., 2019), and interesting recent studies even showed that deletion of AMPs can improve fly survival after traumatic brain injury (Swanson et al., 2020; van Alphen et al., 2022). An effect of AMP overexpression in aging has also been proposed, albeit with conflicting results. Cao et al. (Cao et al., 2013) reported AMP upregulation and neurodegeneration in the brains of *dnr1* mutant flies, and showed that AMP overexpression in neurons was sufficient to induce neurodegeneration. Badinloo et al. found that chronic and ubiquitous AMP overexpression reduced fly lifespan alongside induction of mitochondrial stress (Badinloo et al., 2018). A recent study of the AMP gene *Metchnikowin* further suggests a trade-off between greater antimicrobial activity and host fitness (Perlmutter et al., 2023). These results suggest AMPs could be deleterious to host fitness, which is also supported by evolutionary studies showing that AMP deletions segregate in wild populations (Early et al., 2017; Hanson et al., 2019b). However in direct contradiction to this idea, an earlier report found that excess expression of certain AMPs could extend lifespan (Loch et al., 2017).

Presently, we do not know how AMPs impact aging, and a direct link between AMPs and aging remains to be demonstrated. Some of the conflicting results mentioned above may arise from use of different conditions (e.g. temperature, sex, mating status, nutrition) (Camus et al., 2019; Dobson et al., 2018; Landis et al., 2015; Miquel et al., 1976), microbiome differences, which could help explain contradictory findings on the impact of germ-free conditions on lifespan (Brummel et al., 2004; Ren et al., 2007), or lack of control over genetic background, including transgene insertions (Ferreiro et al., 2017; Sasaki et al., 2021), all of which can influence longevity. In the absence of mutants, these studies have relied on the use of over-expression or RNAi to modulate AMPs, methodologies with certain limitations (Tower et al., 2017; van der Graaf et al., 2022). As a result, exactly how AMPs contribute to aging remains unclear.

Here we have leveraged a recently generated collection of AMP mutations to analyze whether these immune effectors impact aging. We find that individual AMP deletions do not markedly affect aging, with the possible exception of *Defensin*. However, *ΔAMP14* flies lacking 14 AMP genes displayed reduced lifespan associated with microbial dysbiosis. Rearing *ΔAMP14* flies in germ-free conditions significantly rescues lifespan, indicating that AMPs contribute to lifespan through their impact on the microbiome. In contrast, depleting the microbiome of *Relish* mutant flies did not rescue lifespan, suggesting the Imd pathway can affect lifespan independent of regulating AMP genes. Together, our results indicate that AMPs are likely not direct contributors to the aging process, though they have a major impact on aging through AMP-microbiome interactions. Thus, our study, using loss of function mutations, clarifies the role of these innate immune effectors in aging.

## Results

### Presence of cryptic infections may confound lifespan analyses in *Drosophila*

To address the role of AMP genes in aging, we first compared the lifespan of flies lacking eight AMPs located on Chromosome II (*ΔAMP Chr2* lacking *Def, Dro, AttA,B,C, Mtk*, and *DptA,B)* to their DrosDel isogenic wild-type (*iso w*^*1118*^) controls. Of note, all our mutations are in the DrosDel isogenic genetic background (unless mentioned otherwise), and are negative for the endosymbiont *Wolbachia*. We show the lifespans of male flies in the main text, and female flies in the supplementary figures. In most cases, trends were similar between the two sexes. Cases where trends differed between male and females are noted in the main text. We observed a striking effect where *ΔAMP Chr2* deficient flies displayed a marked lifespan extension compared to wild-types (Fig. 1A). This first result would be consistent with studies suggesting AMPs negatively impact lifespan. However, we were surprised by the short lifespan of our *iso w*^*1118*^ wild-type compared to other aging studies, and to a second wild-type included in these experiments, *Oregon R* (*OR-R* in Fig. 1A). *Drosophila* can carry a number of cryptic bacterial or viral infections that affect fitness (Plus et al., 1976). Notably infection with *Drosophila nora virus* is common in lab stocks, and this virus has been shown to reduce lifespan (Habayeb et al., 2009). We thus suspected some component of the virome/microbiome of our *iso w*^*1118*^ flies could be affecting our *iso w*^*1118*^ wild-type but not *ΔAMP Chr2* flies. We therefore cleared our flies of their virome/microbiome through bleaching, allowed microbiome recolonization by microbes in the food medium, and assessed the lifespan of the cleaned *iso w*^*1118*^ stock (protocol in materials and methods). At the same time, we screened our *iso w*^*1118*^ flies for a panel of common contaminating viruses; specifically *Drosophila sigmavirus, Drosophila A virus, Drosophila C virus*, and *Drosophila nora virus*. We did detect *Drosophila nora virus* (hereafter “*nora*”) in our *iso w*^*1118*^ flies, but not in our *ΔAMP Chr2* mutants, nor in other stocks included in those experiments. We also detected *nora virus* in four other genotypes at different times during our five years of study (*AttC*^*Mi*^, *Bom*^Δ*55C*^, *Group C, OR-R*, defined later). In some cases, these stocks were previously nora-negative, and so, were seemingly contaminated from standard fly tipping (Fig. 1B). Strikingly, the bleaching treatment markedly improved the lifespan of all of these stocks to rival their contemporaries, such as *ΔAMP Chr2* flies. The net lifespan reductions for *iso w*^*1118*^ and *OR-R* were ∼39% and ∼23% respectively, possibly suggesting genetic background-effects in susceptibility to *nora virus*. We conclude that our wild-type reference had artificially reduced lifespan, likely due to a cryptic *nora virus* infection. While we took care to control for genetic background effects, this stock, at the time, was not an appropriate baseline for comparison. These considerations are in line with recommendations that healthy wild-type stocks should live to median lifespans of 70-90 days (Piper and Partridge, 2018), emphasizing the importance of considering the health of our control stocks prior to making comparisons across stocks/genotypes.

**Figure 1:**
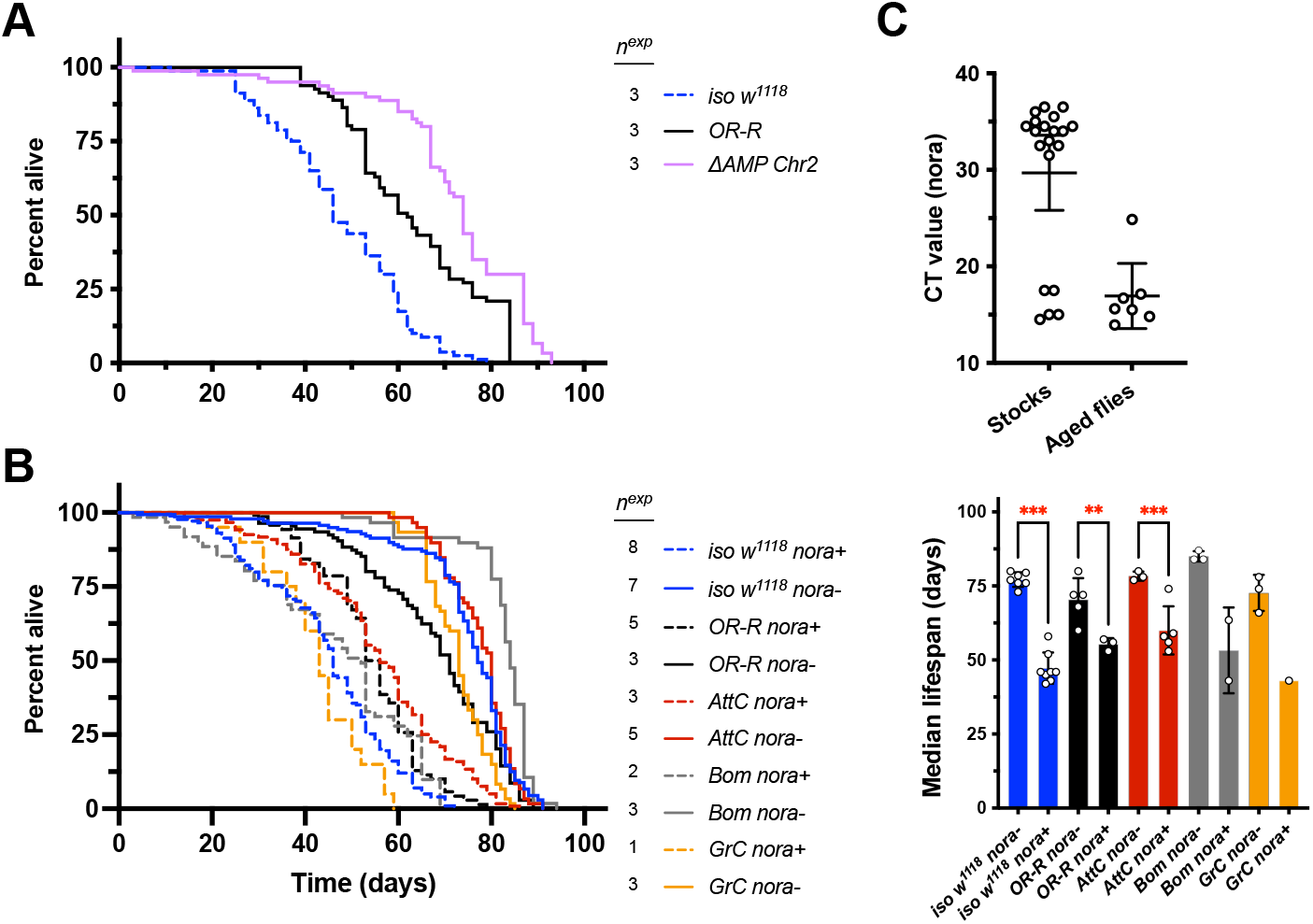
Drosophila nora virus significantly reduces lifespan of DrosDel iso w^1118^ flies. Male flies are reported here and female flies in Fig. S1. A) Comparison of lifespan from early experiments using two wild-types (iso w^1118^ and OR-R) alongside compound AMP mutants lacking Def, Dro, AttA,B,C, Mtk, and DptA,B, which are deleted in ΔAMP Chr2 flies. B) Effect of nora clearance on lifespan of iso w^1118^, OR-R, AttC, Bom, and GrC genotypes. Median lifespans are shown in the right panel (** P < .01, *** P < .001). Number of independent experiments (n^exp^) is reported. C) Nora titres measured by CT value in 18°C source stocks (Stocks) or nora-positive flies aged 3+ weeks kept at densities of 20 flies per vial (Aged flies). CT values represent nora titre from 5ng total fly RNA per 10μL qPCR reaction.

After detecting this contamination in our wild-type flies and seeing the impressive deleterious effect *nora* had on lifespan in our hands, we screened various lab stocks for *nora* virus. We found that 20 out of 44 arbitrarily selected stocks (both from our lab and others) were *nora-*positive, though *nora* titres were typically found only at low levels under standard rearing conditions. We also checked RNA samples collected from aged *nora*-positive flies (aged >3 weeks), finding consistent high *nora* titres (Fig. 1C). These stocks also often showed abdominal bloating at older ages pre-empting mortality (personal observation), similar to bloating seen in flies infected with *Drosophila C virus* (Chtarbanova et al., 2014), or after systemic infection with some strains of *Acetobacter* bacteria (Hanson et al., 2022b). We thus suspect *nora virus* contamination contributed greatly to the early mortality of our *iso w*^*1118*^ flies in our first lifespan experiments (Fig. 1A), with a more drastic effect than previously reported (Habayeb et al., 2009). However, we will note that we did not intentionally perform infection *nora* experiments, and so here we provide only correlation-based evidence.

### Methodology for measuring lifespan and control genotypes

Following these results, we screened all our fly stocks for *nora virus* and bleached *nora*-positive stocks before use in experiments. Subsequent longevity experiments were carried out with the following conditions: we used our standard food medium (recipe in materials and methods), flipped flies three times per week, with ∼20 flies per vial, using mated males or females (sexes kept separate), and performed lifespans at 25°C unless otherwise specified. As an assay for brain health, we also monitored locomotor competence during aging using the climbing pass rate assay at 5, 40, 50, and 60 days post-eclosion (dpe) unless specified otherwise.

To compare lifespan, we first used a Cox mixed model commonly used in the literature. However, we found this statistical method was overly-sensitive for our purposes, as even minor differences in lifespan were returned as highly significant (P < .001). In some cases, this was driven by sex*genotype interaction effects. Because we kept males and females in separate vials, we realized that any putative sex*genotype interactions present in our Cox mixed model were indistinguishable from vial effects. This is especially important as a consideration, as our AMP mutants are known to suffer dysbiosis with aging ((Marra et al., 2021) and see below), compounding the impact of vial-specific microbiome stochasticity (File S4 for further discussion). Visual inspection of survival curves suggests that, by and large, even significant Cox mixed model variation in survival largely reflects variation around the mean wild-type lifespan, which is expected when performing multiple hypothesis tests (Streiner and Norman, 2011). Thus, we paired our lifespan analyses with one-way ANOVA statistics run on median lifespans per experiment, intended to get a more stringent measure of lifespan differences by placing greater value on effects that were consistent across experiments. As such, we use one-way ANOVA *P*-values by default in the text to report significant lifespan differences.

Before we address the impact of AMP mutations on lifespan, we analyzed the lifespans of mutants affected in the Toll (*iso spz*^*rm7*^) or Imd (*iso Rel*^*E20*^) pathways that have been backcrossed in the DrosDel background (Ferreira et al., 2014). We also compared our DrosDel *iso w*^*1118*^ *white*^-/-^ flies with another wild-type (*OR-R*), which has *white*^*+/+*^ red eyes. Of note, *white* gene deletion may affect lifespan through e.g. the role of *white* in the brain (Ferreiro et al., 2017), or through *white* regulation of intestinal stem cell proliferation (Sasaki et al., 2021). We further included non-isogenic flies with previously-reported lifespan effects to show how control genotypes behave in our conditions, and set expectations for the size and consistency of lifespan effects we could reasonably observe in our hands compared to other studies. This included: *methuselah* mutants suggested to have exceptionally long lifespan (*mth*^*1*^) (Lin et al., 1998), a *dnr1* mutation that reduces lifespan previously associated with neurodegeneration and aberrant AMP induction (*dnr1*^*2-133*^) (Cao et al., 2013), and *ATM*^*8*^ flies with precocious neurodegeneration caused by autophagy defects, also associated with Imd pathway activation (Petersen et al., 2013); *mth*^*1*^, *dnr1*^*2-133*^, and *ATM*^*8*^ were not backcrossed into the DrosDel background, and have red eyes. Our experiments confirmed that *ATM*^*8*^ flies have significantly shorter lifespan (Fig. 2A,C) associated with poor climbing competence (Fig. 2D). We also observed that our *OR-R* wild type displays a shorter lifespan compared to our DrosDel *iso w*^*1118*^ wild-type. Contrary to expectation, the lifespan of *mth*^*1*^ flies was not longer than wild-type (P > .10, Fig. 2A,C). However, we did observe improved climbing competence of *mth*^*1*^ flies into old age (Fig. 2D), which suggests these flies do have a form of improved fitness with aging in our hands. These experiments also confirmed a reduced lifespan in *Rel*^*E20*^ flies as found by other studies (Kounatidis et al., 2017; Petersen et al., 2013), suggesting a significant contribution of the Imd pathway transcription factor *Relish* to aging.

**Figure 2:**
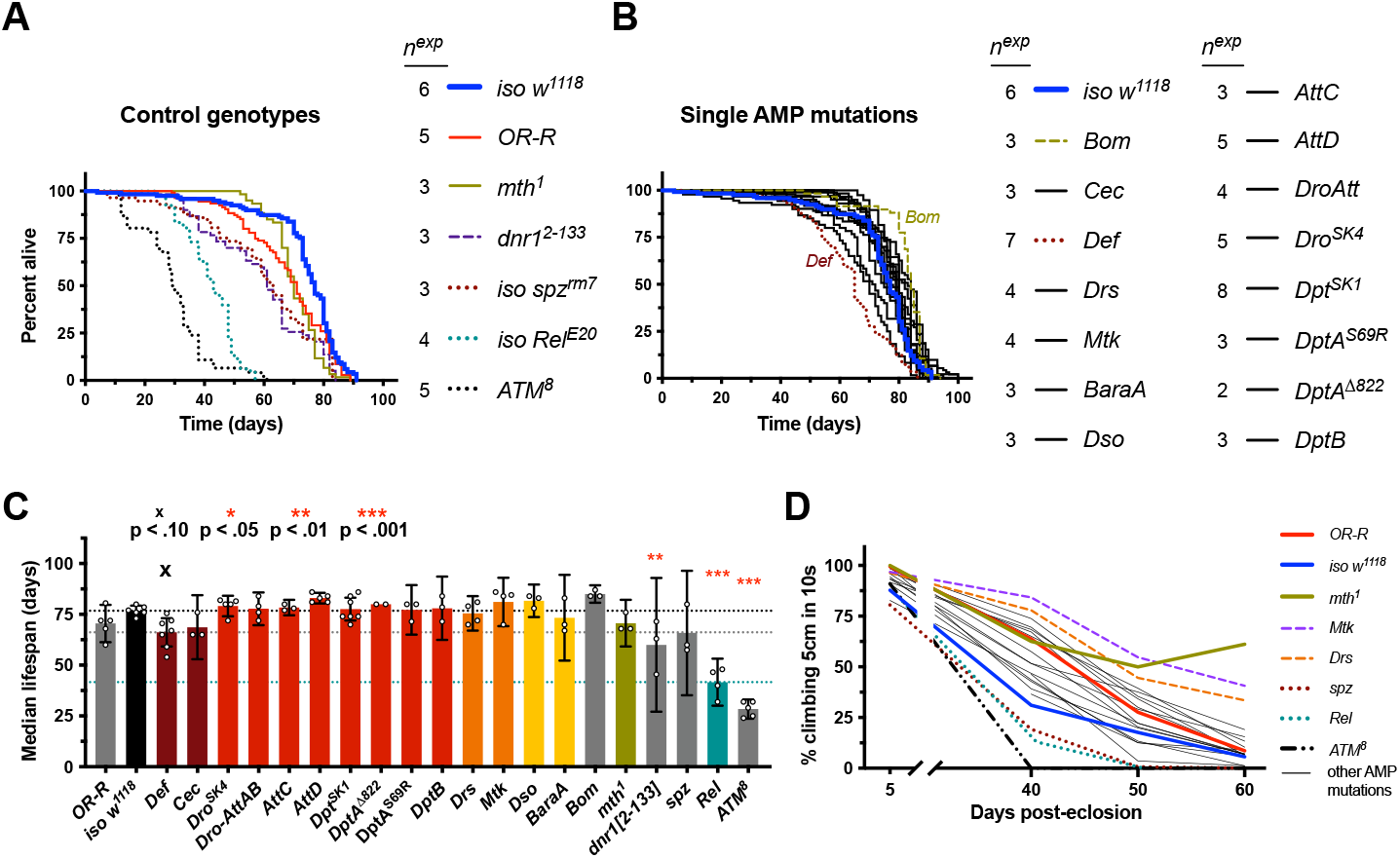
Individual AMP gene deletions do not drastically affect lifespan. Male flies are reported here and female flies in Fig. S3. A) Cumulative lifespans of flies with various genetic backgrounds. Of note, ATM^8^ data are based on fewer individuals per experiment (see File S1). B) Cumulative lifespans of single gene AMP mutants. Most AMP mutant lifespans (black lines) cluster around the wild-type (blue line), except Def^SK3^. Bom^Δ55C^ is also noted as an outlier perhaps living slightly longer than iso w^1118^, which was not seen in females (Fig. S3). C) Median lifespans where each data point represents one replicate experiment (cumulative of ∼20 males). Median lifespan analysis suggests that the only AMP mutation noticeably differing from iso w^1118^ was Def^SK3^. Of note, the impact of Def on lifespan was not corroborated using RNAi (Fig. S4). Bom^Δ55C^ median lifespans were not different from iso w^1118^. Horizontal dotted lines indicate median lifespans of iso w^1118^ (top), Def^SK3^ (middle), or Rel^E20^ (bottom). Statistic summaries (p-values: ^**x**^, *, **, ***) reflect comparisons to iso w^1118^. D) Climbing pass rates suggest most AMP mutants climb like wild-type flies, while methuselah mutants uniquely retain climbing competence into old age (also seen at 29°C, Fig. S2); Mtk and Drs males also show improved climbing with aging (but see supplemental text in File S4).

We also repeated these lifespan experiments at 29°C, which represents a more stressful temperature for *Drosophila melanogaster*, causing more rapid aging (Miquel et al., 1976). When we reared flies at 29°C, as expected, we observed precocious aging both in terms of lifespan and climbing pass rate compared to 25°C for all genotypes tested: *iso w*^*1118*^, *OR-R, dnr1*^*2-133*^, *mth*^*1*^, and *Rel*^*E20*^ (Fig. S2A,B). Again, *mth*^*1*^ flies retained climbing competence longer than other genotypes (Fig. S2C), despite no lifespan extension effect. Of note, differences in lifespan between *OR-R* and *iso w*^*1118*^ were lost when flies were raised at 29°C. The previous *dnr1* mutant study showing reduced lifespan used 29°C (Cao et al., 2013), and *dnr1*^*2-133*^ generally had reduced lifespan compared to *iso w*^*1118*^ and *OR-R* wild-types at 29°C, albeit only significantly comparing female *OR-R* and *dnr1*^*2-133*^ flies (P < .05), while 25°C experiments were not significant compared to *OR-R* (P > .10).

Collectively, we observed that, *ATM, dnr1*, and *Relish* deletion reduced lifespan consistent with previous studies (Cao et al., 2013; Kounatidis et al., 2017; Petersen et al., 2013; Shaposhnikov et al., 2014). Surprisingly, we did not observe lifespan extension in *methuselah* mutants. However, *mth*^*1*^ flies retain climbing competence far better than any other genotype assayed, indicating these flies do have a form of improved aging in our hands. Thus, we could broadly replicate the trends of findings from previous studies, though lifespan itself was not always repeatable in our hands.

### Single AMP gene deletion does not drastically affect lifespan

We next analysed the lifespan of flies lacking individual or small genomic clusters of AMP genes at 25°C. This included mutants for all ‘classical’ AMPs from the seven gene families initially identified: *Def, Cec* (removing 4 *Cecropins*), *Dro* alone, and also *Dro* alongside *AttA* and *AttB*, and then *AttC, AttD, Mtk, Drs, DptA* and *DptB* together or individually, as well as mutants for the more recently described Toll-regulated effector genes *Daisho (Dso1* and *Dso2), Baramicin A (BaraA)*, and *Bomanins (Bom*^*Δ55C*^, removing ten *Bom* genes at cytogenetic map 55C).

We found no major effect on lifespan of any AMP mutation individually, as survival curves tended to disperse randomly around the *iso w*^*1118*^ lifespan curve, and median lifespans were not significantly different from *iso w*^*1118*^ (Fig. 2B-C). The only exception to this trend was male *Def*^*SK3*^ flies, which had noticeably reduced lifespan compared to *iso w*^*1118*^ (P = .059). In general, climbing competence was also distributed in a wild-type range comparable to *iso w*^*1118*^ or *OR-R* flies, and most were not exceptionally good climbers into old age like *mth*^*1*^ flies, though *Mtk* and *Drs* males had somewhat improved climbing into old age (Fig. 2D, Fig. S5). The *Def*^*SK3*^ reduction in lifespan was sufficiently interesting that we tested this effect using ubiquitous (*Actin5C-Gal4*) or glia-specific (*Repo-Gal4*) *Def* interfering RNA (*Def-IR*, RNAi), however we saw, if anything, the opposite effect by silencing *Def* by RNAi compared to the results seen with the *Def*^*SK3*^ mutant (*Act>Def-IR* males had longer lifespans, and females shorter lifespans). However, RNAi genetic controls also suggested complex genetic background effects independent of the Gal4 and RNAi constructs, confounding meaningful interpretations of those results. In general, we could not support a reduced lifespan effect of single AMP mutation with RNAi (Fig. S4).

Collectively, our study using isogenic mutants shows that individual AMP mutations do not significantly affect lifespan, with the possible exception of *Def*^*SK3*^. Overall, we conclude that deleting single AMP genes has no effect on lifespan beyond levels of difference we also observed when comparing different wild-types.

### ΔAMP14 flies lacking seven AMP gene families display significantly reduced life span

*Drosophila* possess many AMPs with possibly redundant or synergistic activities. Thus, AMPs could affect lifespan only when several genes are deleted simultaneously. After evaluating the effect on lifespan of individual AMP genes or genomic clusters of AMPs, we investigated whether combinatory loss of AMPs could impact aging. Using the same approach as described in Hanson et al. (Hanson et al., 2019a), we generated four groups of compound AMP mutants that remove different subsets of AMP gene families: *Group A* flies were deleted for *Defensin* and the four *Cecropins. Group B* flies were deleted for the structurally related *Drosocin, Attacin*, and *Diptericin* families. *Group C* flies were deleted for the two antifungal *Drosomycin* and *Metchnikowin* peptides. And we introduce a new isogenic line combining the loss of two recently-described Toll-regulated antifungal peptides: *“Group D”* that are deleted for both *Baramicin A* (Hanson et al., 2021) and the two *Daisho* genes (Cohen et al., 2020). We screened combined mutants lacking each of these AMP groups, including all the combinations of Groups *A, B*, and *C* (i.e. *AB, AC, BC*, and *ABC* (aka *ΔAMP14*)). We also screened flies lacking 10 AMP genes, but which retain a wild-type *Cecropin* locus (*ΔAMP10*, as used previously (Carboni et al., 2021; Hanson et al., 2019a)). In total we screened nine AMP combinatory genotypes for lifespan and climbing effects (Fig. 3).

**Figure 3:**
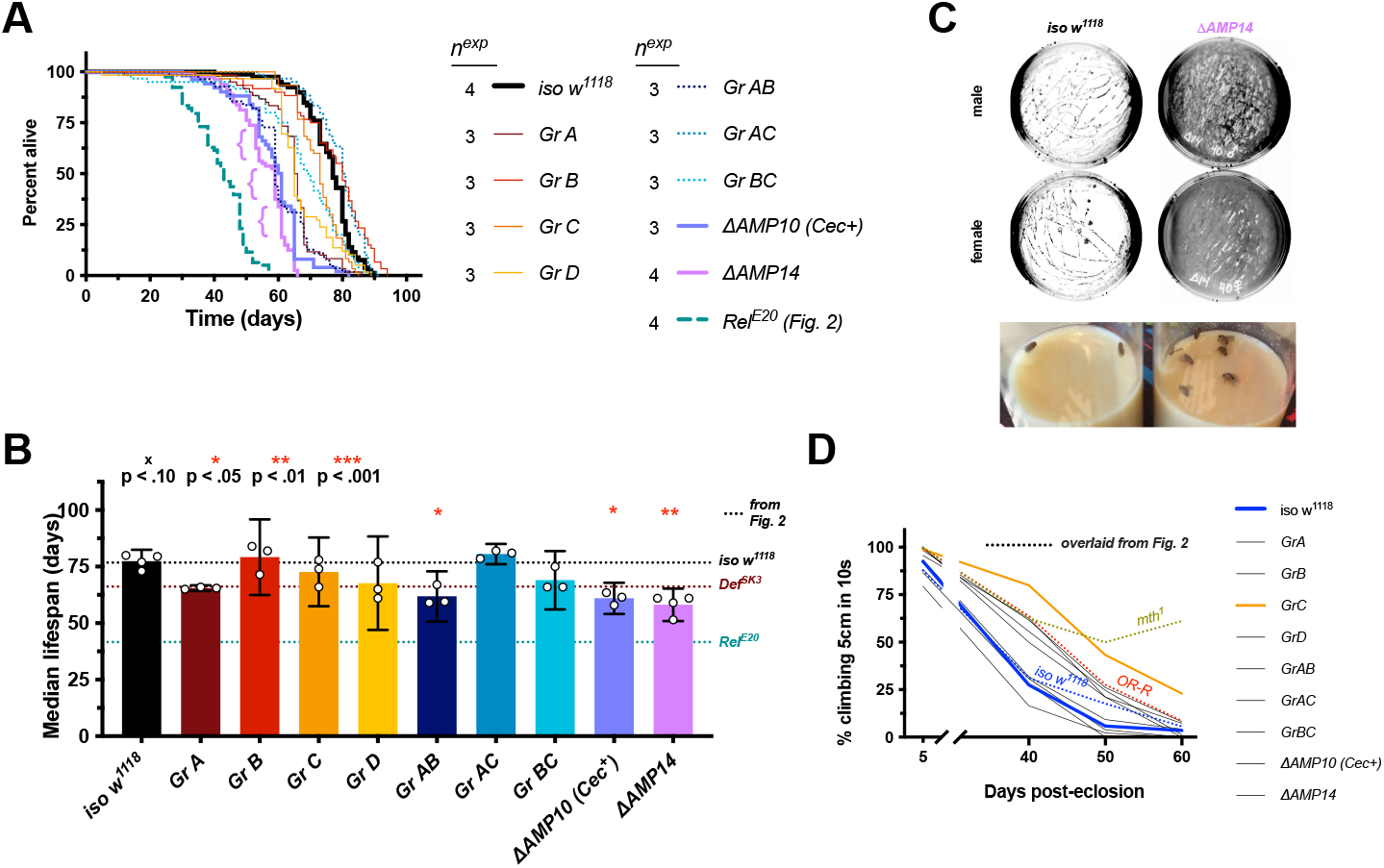
ΔAMP14 flies have significantly reduced lifespan. Male flies are reported here and female flies in Fig. S6. A) survival curves of various compound AMP mutants. The lifespan of Rel^E20^ from Fig. 2 is overlaid for direct comparison. “{” annotations highlight major mortality events in ΔAMP14 flies. B) Median lifespans of compound AMP mutants. Dotted lines indicate average median lifespans from Figure 2 of iso w^1118^ (top), Def^SK3^ alone (middle), and Rel^E20^ (bottom) for easier comparisons across figures. Statistic summaries (p-value: ^**x**^, *, **, ***) reflect comparisons to iso w^1118^ data specific to Figure 3. C) Top: representative photonegative images of agar plates seeded by the microbiome found in the vial of 40-day old flies. Thick bacterial films in ΔAMP14 vials are readily visualized by this method, which shows the significantly greater bacterial density (dark parts) compared to iso w^1118^. Bottom: representative photo of iso w^1118^ and ΔAMP14 food vials revealing discoloured bacterial biofilm alongside a major mortality event (“{“ in Fig. 3A). D) Climbing pass rates of AMP group mutants, with climbing curves from genotypes in Figure 2 overlaid for direction comparison. Group C is highlighted for having a slightly improved climbing over aging, though this improvement is still minor compared to the climbing competence of mth^1^ flies (also see females in Fig. S6 and File S4).

We found a significant effect of *Group A* mutations on male lifespan compared to *iso w*^*1118*^, which is not unexpected given this AMP group uses the same 2^nd^ chromosome as the isogenic *Def*^*SK3*^ mutation as above (Fig. 2B), except with the additional loss of *Cecropins* on the third chromosome. Comparisons between the two suggest a non-significant effect of *Cecropin* mutation (*Def*^*SK3*^ vs. *Group A*, P > .10). Deleting other groups of AMPs typically resulted in non-significant effects on median lifespan compared to their *iso w*^*1118*^ control (P > .10). *Group AB* flies also had reduced lifespan compared to *iso w*^*1118*^, but were comparable to *Group A* alone (*Group A* vs. *Group AB*, P > .10). *Group AC* and *BC* flies had comparable lifespans to *iso w*^*1118*^ (P > .10 in both cases), which is notable as *Group AC* flies have the *Def*^*SK3*^ mutation, but only after an additional round of chromosome II recombination with the isogenic *Mtk*^*R1*^ chromosome needed to generate the *Group AC* genotype. Deleting ten AMP genes, which included *Defensin*, also had a *Group A*-like lifespan (*ΔAMP10* vs. *Group A*, P > .10). Thus, various combinations of mutants had lifespans comparable to either *iso w*^*1118*^ or *Def*^*SK3*^ alone. However, deletion of all 14 classic *Drosophila* AMP genes caused a more pronounced median lifespan reduction than the effect of *Group A* alone (*ΔAMP14* vs. *Group A*, male P = .059, female P = .002). There was minor variation in climbing competence amongst AMP mutant groups, with *Group C* flies (*Mtk, Drs* double mutants) having slightly improved climbing into old age. However, this effect was minor compared to the climbing competence of *mth*^*1*^ flies into old age (Fig. 3D and see supplementary text).

While deletion of any single AMP gene had little impact on lifespan, deletion of all 14 classical AMP genes causes lifespan reduction significantly different from wild-type flies. Moreover, recombination of *Def*^*SK3*^ with another mutation to produce *Group AC* flies yields lifespan comparable to *iso w*^*1118*^. This result reinforces the need for careful interpretation of the *Def*^*SK3*^ mutation effect of *Group A*. Despite this caveat, *ΔAMP14* flies had significantly reduced lifespan compared to both wild-type flies and other AMP groups. The lifespan of *ΔAMP14* flies is intermediate between *iso w*^*1118*^ and *Rel*^*E20*^, indicating *Relish* likely impacts fly lifespan through processes other than AMP regulation.

### *ΔAMP14* flies suffer microbial dysbiosis with aging

During our studies, we noticed that the food surface of the vials containing *ΔAMP14* flies became discolored and sticky upon aging, suggesting microbial proliferation (Fig. 3C bottom). The *Drosophila* microbiome is found both in the gut and in the external environment due to constant ingestion and fecal deposition (Broderick et al., 2014; Pais et al., 2018; Storelli et al., 2018). We suspected this change in the fly food appearance in *ΔAMP14* upon aging could be linked to a change in the microbiome, as we have recently described a role for AMPs in regulating *Acetobacter* using *ΔAMP14* flies (Marra et al., 2021). Sticky and discolored food was also observed in *Rel*^*E20*^ flies, which do not express antimicrobial peptides, and indeed both *ΔAMP14* and *Rel*^*E20*^ flies suffer increased dysbiosis over aging (Marra et al., 2021). To test if microbiome load was associated with early mortality of *ΔAMP14*, we monitored bacterial abundance on the food medium. To do this, we emptied vials where flies had been present for two days, added glass beads to those vials, shook vials with beads for 10 seconds, then placed those beads on agar plates and spread vial microbes by rolling the beads over the agar for 10 seconds. By 40 days post-eclosion (dpe), plating of vial contents using beads confirmed far higher microbe loads in aged *ΔAMP14* vials compared to *iso w*^*1118*^ (Fig. 3C top). Moreover, in the time period between 50-70dpe, most major mortality events in *ΔAMP14* vials were associated with sticky and discolored bacterial food (example photo in Fig. 3C bottom). These major mortality events are also seen as precipitous drops in Fig. 3A (indicated with “{“) survival curves in *ΔAMP14* flies, and similar trends were observed for *ΔAMP10*.

We conclude that *ΔAMP14* flies thus suffered increased microbe load within vials with aging, agreeing with an increase in gut microbiome abundance with aging shown previously using gnotobiotic flies (Marra et al., 2021). This suggests that precocious aging observed in *ΔAMP14* flies could be indirectly caused by the impact of AMPs on the microbiome.

### Lifespan of *ΔAMP14* flies can be rescued significantly by rearing on antibiotic medium

To test whether the effect of AMP mutation on lifespan was linked to changes in the microbiome, we performed the same lifespan experiments in germ-free conditions using *iso w*^*1118*^, *ΔAMP14*, and *Rel*^*E20*^. For this, we first bleached embryos, and then kept larvae and emerging adults on antibiotic food media for their entire lifespan. In our hands, antibiotic-reared *iso w*^*1118*^ flies have a similar lifespan compared to conventionally reared flies. Likewise, in our hands *Rel*^*E20*^ mutants had similar lifespan whether reared conventionally or in antibiotic conditions. However, rearing *ΔAMP14* flies on antibiotics significantly rescued their lifespan (Fig. 4A-B, *ΔAMP14* ABX vs. CR, P = .013), and a similar non-significant trend was seen in females (Fig. S7A-B). Climbing competence remained largely unchanged, although male Δ*AMP14* flies showed improved climbing specifically at 50dpe in germ-free conditions (P = .053, Fig. 4C).

**Figure 4:**
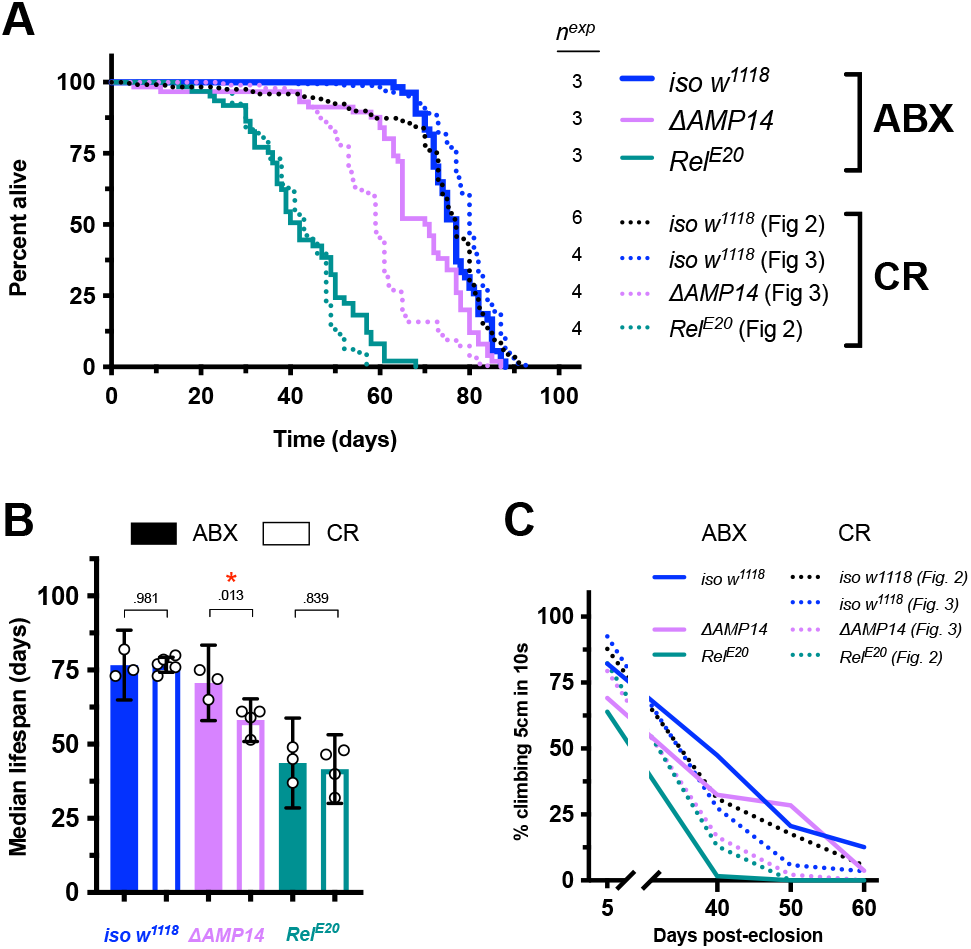
Microbiome depletion rescues the ΔAMP14 fly lifespan. Male flies are reported here and female flies in Fig. S7. A) Survival curves, including both antibiotic-reared flies (ABX), and also conventionally-reared (CR) lifespans from previous figures as dotted lines for direct comparison. B) Median lifespans, including both ABX and CR fly lifespans for direct comparison. (conventionally-reared iso w^1118^ and Rel^E20^ lifespans shown in Fig. 2C). C) Climbing pass rates of ABX (solid lines) and CR (dotted lines) flies at 5, 40, 50, and 60 days post-eclosion.

These results suggest that the short lifespan of *ΔAMP14* flies primarily relies on the impact that antimicrobial peptides have on the microbiome through aging. The observation that antibiotics treatment rescues *ΔAMP14* but not *Relish* mutant lifespan suggests the short lifespan of *Rel*^*E20*^ flies depends on effects other than the microbiome and independent of AMPs.

## Discussion

A number of studies done in *Drosophila* and *C. elegans*, but also vertebrates, have implicated AMPs in processes as diverse as behavior, neurodegeneration, tumor clearance, and aging. In mammals, some AMPs function as pro-inflammatory cytokines, and as such could influence these processes by disrupting homeostasis when chronically expressed (Liang and Diana, 2020; Mookherjee et al., 2020; Smith and Gwyer Findlay, 2022). However, to date there is no evidence of a cytokine role for AMPs in *Drosophila* (Hanson and Lemaitre, 2020). AMPs can also disrupt the membranes of aberrant host cells, that may become more negatively charged due to the exposure of phospholipids such as phosphatidylserine, a well-known “eat-me” signal that promotes the phagocytosis of apoptotic cells (Fadok et al., 1992; Hakim-Mishnaevski et al., 2019; Manaka et al., 2004). Studies have now also implicated AMPs in the control of tumorous growth in vivo in *Drosophila* (Araki et al., 2019; Parvy et al., 2019), providing a proof of principle that AMP action can target self-tissue. Thus, AMPs could have a role in tissue maintenance with consequences on the aging process. Moreover, AMPs have a number of properties in common with neuropeptides, often being cationic and amphipathic (Augustin et al., 2017; Smith and Gwyer Findlay, 2022), alongside descriptions of AMP-like genes undergoing downregulation in the brain after exposure to pheromones (Gendron et al., 2014), modulating sleep (Sinner et al., 2021; Toda et al., 2019), memory (Barajas-azpeleta et al., 2018), and affecting behaviors (Ebrahim et al., 2021; Hanson et al., 2021; Kobler et al., 2020). Thus, it cannot be excluded that AMPs modulate neuronal activity, a process that could impact lifespan.

There is an increase of AMP gene expression observed upon aging, though it was unclear if AMPs directly affect lifespan, or if this activation is just a secondary consequence of aging. For instance, aging is accompanied by dysbiosis and gut barrier dysfunction allowing opportunistic systemic infection by microbiome bacteria (Buchon et al., 2009; Clark et al., 2015; Liu et al., 2022; Marra et al., 2021; Rera et al., 2012), which should activate immune signalling and induce AMP expression. Using flies carrying null mutations in AMP genes, we find no evidence that individual AMPs are so essential to host physiology that they have a notable impact on fly lifespan. Overall, the lifespan of flies mutated for individual genes or clusters of AMPs, *Bomanins*, or *Daishos*, did not differ from the wild-type. The climbing activity of these mutants were also similar to wild-type flies. The only possible exception to this was the apparent lower lifespan of *Def* mutant males. However conflicting results from RNAi experiments and use of *Group AC* flies with the *Def*^*SK3*^ mutation after additional rounds of recombination suggest the somewhat shortened lifespan of *Def* mutant males was not linked to *Def* mutation itself, but due to the presence of one or several cryptic mutations that were not removed during the isogenization process due to their proximity to the *Def* locus. Alternately, complex interactions amongst AMPs could provide a protective effect against *Defensin* mutation, as AMP interactions can synergize to prevent damage to host membranes (Drab and Sugihara, 2020). Overall, our study suggests that individual AMPs do not affect lifespan beyond levels of difference we also saw when comparing wild-types and/or classic aging mutant flies (e.g. *mth*^*1*^), a result that contradicts other studies using RNAi or overexpression methods. However, our results do support a role for AMPs in regulating the microbiome over aging. Indeed, we recently showed that *Acetobacter* microbiome bacteria grow out of control in the microbiome of Δ*AMP14* flies (Marra et al., 2021), and later confirmed that *Diptericin B* has a highly specific and important role in suppressing *Acetobacter* growth after systemic infection, which causes bloating similar to what we saw in *nora virus*-infected flies (Hanson et al., 2022b). This phenotype of *Acetobacter* systemic infection could help explain why flies bloat upon infection by enteric pathogens (like *nora virus* or *Drosophila C virus*), or upon aging, given the eventual invasion of gut microbes into the hemolymph (Clark et al., 2015; Rera et al., 2012).

We cannot exclude that additional AMP mutant combinations could reveal a stronger impact of AMPs on lifespan. Indeed, a number of immune-induced peptides that could be AMPs await description (Hanson and Lemaitre, 2020; Schlamp et al., 2021), and additional deletion of those peptides could explain the lifespan difference between *ΔAMP14* and *Relish* mutants. Use of other rearing conditions could also reveal a role for AMPs not found here. As part of this study, we also monitored AMP expression upon aging in four wild-type backgrounds (a lab standard *w*^*1118*^, *Exelexis, OR-R*, and *Canton-S*) by separating out dissected heads and bodies (thorax + abdomen). In our hands, we observed increased expression of AMP genes in fly bodies aged 40 days compared to young flies, though the extent differed by genotype (Table S1 and File S2). However, we saw no marked AMP increase in the head with aging, including in a separate experiment following flies over multiple time points (File S2). Indeed, when AMPs were upregulated in the head upon aging, this was often coupled with a stronger upregulation in the body, suggesting that this increase of AMP expression in the head could derive from head-specific fat body responding like the fat body in the thorax and abdomen. We also tested if glia-specific *Relish* knockdown (via *Repo-Gal4* and *UAS-Rel-IR*) could rescue lifespan, as many studies suggest a role of *Relish* in neurons/glia to rescue neurodegenerative syndromes (Cao et al., 2013; Kounatidis et al., 2017; Petersen et al., 2013; Shukla et al., 2019), but if anything we saw the opposite effect: female *Repo>Rel-IR* flies had shorter lifespan than genetic controls, and we found that *Repo-Gal4* alone had improved climbing into old age, suggesting a genetic background effect on aging unrelated to RNAi using this driver (Fig. S8). While *Repo>Rel-IR* can rescue various models of neurodegeneration, taken together our results suggest AMPs are not especially upregulated in the head upon aging (except when also upregulated in the body), and that glia-expressed *Relish* does not have a major deleterious role in lifespan using our standard rearing conditions. Differences between research groups (local microbiome composition, including cryptic viral infections, food recipe, etc…) could account for these conflicting results. As we found by including *methuselah* mutant flies, improvements to healthy aging could also be consistent across research groups, but actual lifespan extension may be lab-specific. These factors, and the importance of assaying multiple healthy aging metrics, should be considered when comparing our AMP mutant results to the larger field.

Importantly, our study suggests that AMPs collectively affect fly lifespan through their impact on the microbiome. Several studies have investigated the role of the gut microbiome on lifespan using germ-free conditions with mixed results (Brummel et al., 2004; Cai et al., 2021; Iatsenko et al., 2018; Ren et al., 2007; Shukla et al., 2021). It could be that discrepancies between studies reporting an impact of the microbiome on aging results from a complex interaction between nutrition and gut bacteria. In poor or unbalanced diets, the microbiome could have a positive impact on lifespan by extracting more nutrients from the food (Camus et al., 2019; Chaston et al., 2016; Consuegra et al., 2020; Erkosar et al., 2017; Newell and Douglas, 2014; Storelli et al., 2018; Téfit et al., 2018). In contrast, on a lab standard nutrient-rich diet, as we have used in this study, the microbiome could have less impact. Consistent with this, we did not see major differences between germ-free and conventionally raised wild-type flies in this study. Previous studies have already revealed the role of Imd signalling in controlling microbiome load and diversity, preventing dysbiosis (Guo et al., 2014; Li et al., 2016; Liu et al., 2022; Yamashita et al., 2021). The specific microbiome of a given research group could also change the impact of germ-free conditions. For instance, different *Acetobacter* strains have different virulence to the fly during systemic infection (Hanson et al., 2022b), which accompanies aging and intestinal barrier dysfunction (Clark et al., 2015; Rera et al., 2012). Indeed, use of *ΔAMP14* flies revealed a key role of AMPs to control *Acetobacter* levels in the microbiome, and accordingly, *ΔAMP14* flies have increased *Acetobacter* loads upon aging (Marra et al., 2021).

Our present study shows that the action of AMPs preserves lifespan, and that this effect is largely due to their impact on the microbiome. Thus, the impact of AMPs on lifespan is consistent with their well-established microbicidal activity. Interestingly, raising *Relish* mutant flies in axenic conditions did not lead to lifespan extension, indicating that *Relish*, and likely the Imd pathway, have a much more profound impact on host physiology independent of AMP regulation. In line with this, *Relish* and the Imd pathway have been implicated in neural systems (Kounatidis et al., 2017; Masuzzo et al., 2019; Shukla et al., 2019), cell competition (Meyer et al., 2014; Nandy et al., 2018), metabolism (Molaei et al., 2019; Musselman et al., 2018), and enterocyte delamination (Liu et al., 2022; Zhai et al., 2018), processes that likely impact lifespan. It is interesting to note that the food media of *ΔAMP14* and *Rel*^*E20*^ flies was enriched in bacteria, agreeing with elevated bacterial loads in these flies with aging shown previously (Marra et al., 2021). External microbes colonize the *Drosophila* gut, and gut microbes are released into the external environment as part of excreta (Broderick et al., 2014; Newell and Douglas, 2014; Storelli et al., 2018; Winans et al., 2017). Thus, it was expected that an increased gut microbiome load in the absence of AMPs would result in high bacterial load in the fly food medium. Although we reared flies in a vial in artificial lab conditions here, it is tempting to speculate that AMPs expressed in the gut could not only shape the gut microbiome, but also environmental bacteria. The increased load in the food medium could therefore rely either on AMP-mediated control of the gut microbiome, or on external AMPs secreted into the food medium. This would suggest that AMP mutation can exacerbate microbiome effects from both within the fly and in the external vial environment.

During our study, we experienced a number of challenges to lifespan data interpretation. Notably, our reference wild-type and several other fly strains were infected with *Drosophila nora virus*, which had a greater deleterious effect on lifespan of *iso w*^*1118*^ compared to wild-type flies used in a previous study (Habayeb et al., 2009). Thus, we initially interpreted our early results as if AMP deletion extended lifespan to a great extent compared to their isogenic wild-type controls (Fig. 1A). However, our ultimate findings, including standard genotypes and different conditions, instead highlight that loss of AMPs does not extend lifespan; if anything, they show the opposite. We also note that cryptic and chronic infections common in fly stocks, such as *nora virus*, represent a serious threat to aging studies. In our study, we realized this cryptic viral infection confounded our results when comparing our isogenic wild-type lifespan to the expected absolute lifespan of *Drosophila* according to previous recommendations (Piper and Partridge, 2018). We publish this experience, which confused years of data collection, as a cautionary note for the field of aging and immunity. Our hope is that our experience can help others avoid similar confounding factors.

In conclusion, our study reveals a key role of AMPs in the aging process: but mainly indirectly, through their effect on the microbiome. We cannot exclude that certain contexts could reveal an intrinsic effect of AMPs on host tissues during aging, such as conditions found in individuals that have aging-associated diseases like cancer or precocious neurodegeneration, uncommon in standard wild-types. However, here we do not find evidence of AMPs directly impacting aging in a striking way. We are still far from understanding the complex relationship between the immune system, senescence, and aging, which requires further investigation.

## Acknowledgements

We would like to thank Samuel Rommelaere, Jean-Philippe Boquete, Emi Nagoshi, Lukas Neukomm, Kausik Si, and Anzer Khan for helpful discussion. We would also like to thank Brian McCabe, Mariann Bienz, Barry Ganetzky, Steven Wasserman and Lianne Cohen, the Vienna Drosophila Resource centre, and the Bloomington Drosophila Stock Centre for fly stocks requested over the course of this research. This research was supported by Sinergia grant CRSII5_186397 and Novartis Foundation 532114 awarded to Bruno Lemaitre.

## Materials and Methods

### Drosophila rearing conditions

*Drosophila* stocks used in this study, including genotype descriptions, are listed in File S3. Food media used the following recipe (per 600mL): 3.72g agar, 35.28g cornmeal, 35.28g yeast extract, 36mL grape juice, 2.9mL propionic acid, and 15.9mL Moldex. Antibiotic media also contained final concentrations of 50ug/mL Ampicillin, 50ug/mL Kanamycin, 10ug/mL Tetracycline, and 10ug/mL Erythromycin. Flies were flipped three times per week (Monday, Wednesday, Friday), and vials were left on their side to ameliorate the effect of the food medium stickiness on mortality with aging, by allowing flies falling to the ground to drop onto plastic rather than the food surface. This precaution was taken as vial conditions differ markedly in specific immune-deficient genotypes (Marra et al., 2021).

To clear flies of *nora virus*, embryos were collected from grape juice agar plates, rinsed with distilled water, and left to soak in 3% bleach for 3 minutes. Embryos were then rinsed twice in distilled water for one minute each. This protocol was also used to clear the microbiome of antibiotic-reared flies, whereafter we placed embryos directly on antibiotic medium for germ-free experiments.

### Lifespan experiments

Lifespan experiments were conducted from 2017 to 2022. Flies were allowed to emerge and mate randomly for ∼3 days prior to separating males and females. Then, groups of 20 males or 20 females (mated) were flipped three times per week (Monday, Wednesday, Friday) to measure fly lifespan in 90×15mm polystyrene vials.

We used a Cox proportional hazard (CoxPH) mixed effects model to initially analyze lifespan effects, with experimental replicate and biological sex as interaction terms in R version 3.6.3. In both cases, experimental replicate and sex were significant contributors to the model (P < .001). Our impression from the initial data analysis was that the CoxPH model was overly sensitive to minor variation around the geometric mean lifespan of *iso w*^*1118*^ control flies, exacerbated by the large sample sizes used and the many comparisons performed in our study inflating the likelihood of Type I (false positive) statistical errors. Even if minor differences in lifespan were genuine, their ultimate importance was questionable when compared to other genetic backgrounds (e.g. *OR-R, mth*^*1*^), particularly given variable mutation types and transgene insertions used for different AMP mutations (i.e. point mutation, genomic deficiency, *white*^+^, 3xP3-EGFP, or 3xP3-dsRed). We thus preferred to focus on each experiment as if that population of flies represented a single sampling. We therefore treated our data as if we had e.g. n = 3 per genotype (3 experiments), rather than n = 120 (60 males and 60 females) per genotype across 3 experiments. For this purpose, we decided to use median lifespans as our primary readout. Sex-specific median lifespans were analyzed using one-way ANOVA with Holm-Sidak’s multiple test correction implemented in Prism v9.3.1, or one-way ANOVA with Tukey’s HSD correction in R 3.6.3.

### Climbing pass rates

We paired our lifespan data with the gravotaxic locomotor climbing assay to provide an independent metric of aging (Madabattula et al., 2015). This assay assesses the general locomotor competence of the fly, which is often used as a readout of neurodegeneration, but can also reflect generic aging effects (e.g. muscle weakness). Climbing pass rates were filmed for flies at 5, 40, 50, and 60 days post-eclosion ±2 days, and analyzed later manually. A pass was considered if a fly climbed 5cm within 10s of being tapped to the bottom of the vial. Flies were transferred to a chamber made of two empty vials stacked atop each other to provide ample room to climb upwards without reaching the ceiling or disrupting other flies. Two sets of broken markings were made with permanent marker at 2.5cm and 5cm on the lower vial as reference points, though we ultimately report only 5cm climbing rates given similar trends between the two. At each time point, an initial tapping down was performed to associate the flies with their new environment and encourage climbing behaviour, as we found the first repetition of this experiment often had fewer climbers than subsequent repetitions. After this initial association, three technical repeats of the climbing assay were performed, and final values represent the average of these three technical repeats. All climbing measurements were taken between 2pm-5pm to ensure a consistent measurement timeframe.

This assay can be a measure of both climbing speed and/or readiness of response: for instance, older flies sometimes suffered temporary seizures after being tapped down, or might climb slower, more erratically, or simply to a lesser extent than younger counterparts. In most cases, climbing pass data mirrored lifespan data in terms of relative trends. We thus have no suspicion that AMP mutations, individually or collectively, greatly affect locomotory competence with aging. The only noteworthy exception to mirrored trends between lifespan and climbing was *methuselah* mutant flies, which had wild-type lifespans, but improved climbing competence into older age at both 25°C and 29°C.

For analysis of trends of ΔAMP14 in Fig. 4C, we fit a two-way ANOVA measuring the climbing at 40, 50, and 60 days, with an interaction term for age and germ-free conditions in Prism 9.4.1.

### Microbiome monitoring

We monitored the microbiome of compound AMP mutants and antibiotic-reared flies by checking plated vial food contents on MRS + Mannitol agar, a medium amenable to both Lactobacilli and *Acetobacter*. Specifically, at both ∼20 and ∼40 days we took vials where flies had been present for two days, added 5 glass beads to the vials, shook the glass beads in vials for 10s, then transferred glass beads to MRS+Mannitol agar plates and shook plates for 10s, then removed the beads and left plated microbes to grow overnight at 29°C. Vial microbiome loads were checked on Wednesdays. We chose Wednesdays rather than a precise age (e.g. exactly 20dpe) to ensure that flies entered the plating time point after experiencing a similar regimen of flipping; i.e. in our experiments, we only plated microbes from vials where flies had spent the past five days with ∼3 days in a vial accruing microbes over the weekend (Fri-Mon), and the next two days depositing microbes in the vial that was ultimately measured (Mon-Wed). This design was chosen based on demonstrations that flipping regimen drastically affects microbiome load (Arias-Rojas and Iatsenko, 2022; Blum et al., 2013; Pais et al., 2018). Beyond ∼40 days, comparisons were not equal due to onset of mortality in AMP mutants and associated drops in vial fly density.

In antibiotic experiments, microbes were never detected from overnight growth using this method to monitor vial microbiome loads.

### Gene expression assays

Gene expression was performed using primers listed in File S3 with PowerUP SYBR Green Master Mix, using the PFAFFL method of qPCR quantification with Rp49 as the reference gene (Pfaffl, 2001). RNA was extracted using TRIzol according to manufacturer’s protocol. cDNA was reverse transcribed using Takara Reverse Transcriptase.

Dissections of heads from bodies was performed in ice cold PBS, and tubes containing pools of 20 heads or bodies were kept at -20°C until after TRIzol was added to prevent RNA degradation before sample processing.

## Supplementary figure captions

**Figure S1:**
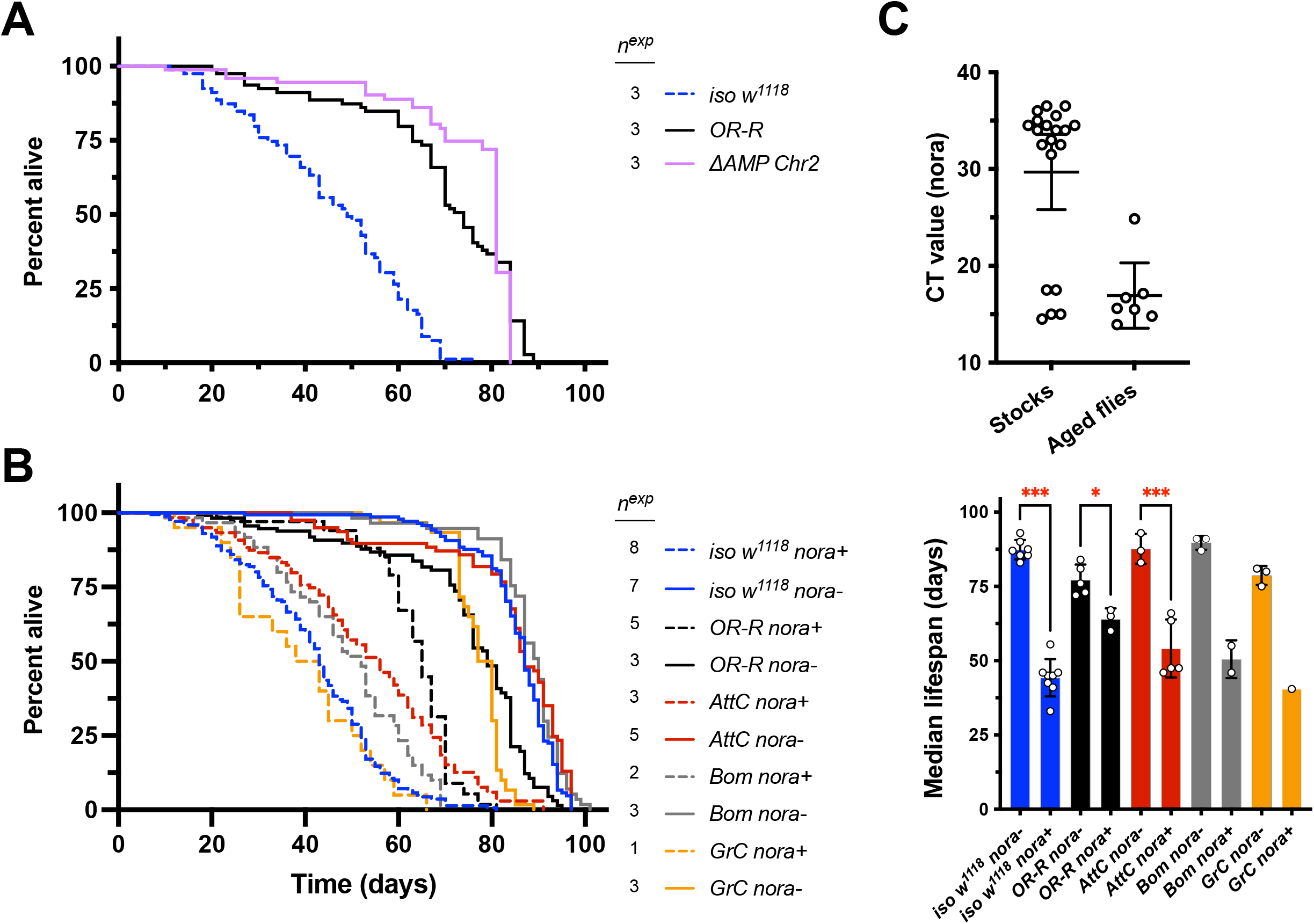
Lifespan of various fly genotypes at 29°C. Male flies are shown in Figure 1. A) Comparison of lifespan from early experiments using two wild-types (*iso w*^*1118*^ and *OR-R*) alongside compound AMP mutants lacking *Def, Dro, AttA,B,C, Mtk*, and *DptA,B*, which are deleted in *ΔAMP Chr2* flies. B) Effect of *nora* clearance on lifespan of *iso w*^*1118*^, *OR-R, AttC, Bom*, and *GrC* genotypes. Median lifespans are shown in the right panel (** P < .01, *** P < .001). Number of independent experiments (*n*^*exp*^) is reported. C) *Nora* titres measured by CT value in 18°C source stocks (Stocks) or *nora*-positive flies aged 3+ weeks kept at densities of 20 flies per vial (Aged flies). CT values represent *nora* titre from 5ng total fly RNA per 10μL qPCR reaction.

**Figure S2:**
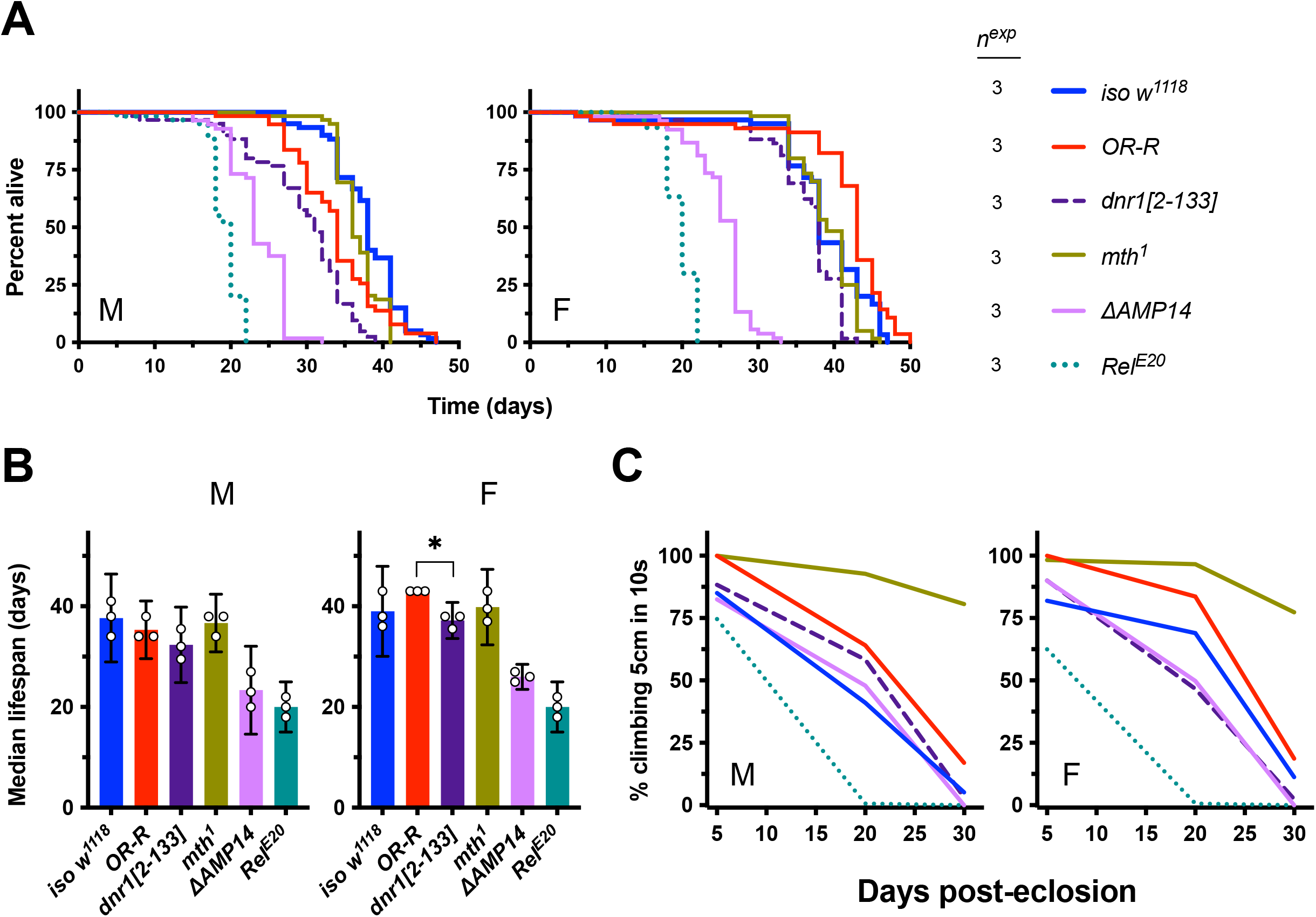
Lifespan of various fly genotypes at 29°C. A) *Relish* mutant (*Rel*^*E20*^) and compound AMP mutants (*ΔAMP14*) suffer significantly reduced lifespan at 29°C. B) Median lifespan data from Figure S2A. C) Climbing pass rates of flies reared at 29°C. *mth*^*1*^ flies are the only fly line that retains climbing competence into old age.

**Figure S3:**
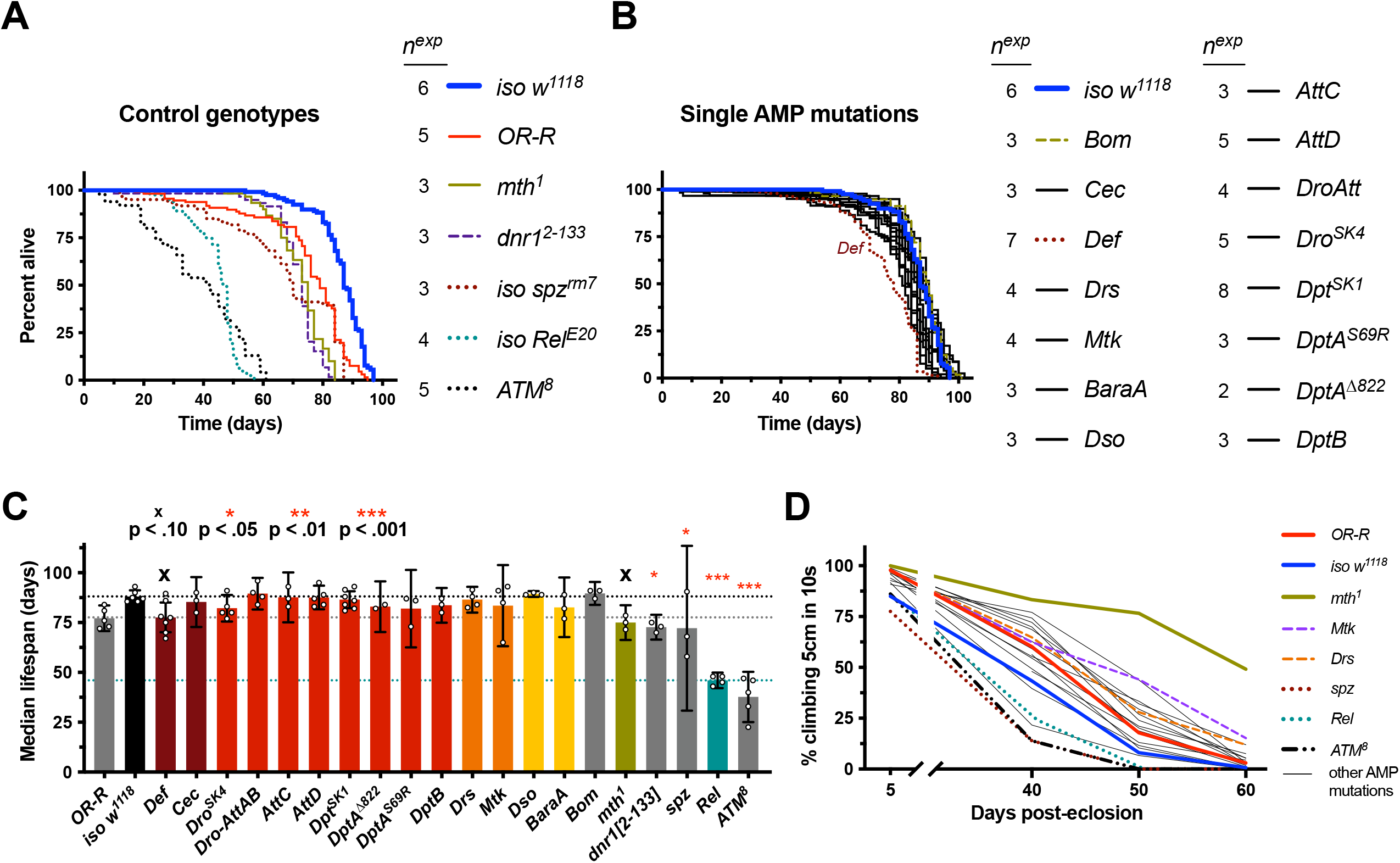
Individual AMP gene deletions do not drastically affect lifespan. Male flies shown in Figure 2. A) Cumulative lifespans of female flies with various genetic backgrounds. Of note, *ATM*^*8*^ data are based on fewer individuals per experiment (see File S1). B) Cumulative lifespans of single gene AMP mutants. Most AMP mutant lifespans (black lines) cluster around the wild-type (blue line), except *Def*^*SK3*^. C) Median lifespans where each data point represents one replicate experiment (cumulative of 20 females). Median lifespan analysis suggests that the only AMP mutation statistically differing from *iso w*^*1118*^ was *Def*^*SK3*^. Of note, the impact of *Def* on lifespan was not corroborated using *Defensin* RNAi (Fig. S4). Horizontal dotted lines indicate median lifespans of *iso w*^*1118*^ (top), *Def*^*SK3*^ (middle), or *Rel*^*E20*^ (bottom). Statistic summaries (p-values: ^**x**^, *, **, ***) reflect comparisons to *iso w*^*1118*^. D) Climbing pass rates suggest most AMP mutants climb like wild-type flies, while *methuselah* mutants uniquely retain climbing competence into old age (also seen at 29°C, Fig. S2).

**Figure S4:**
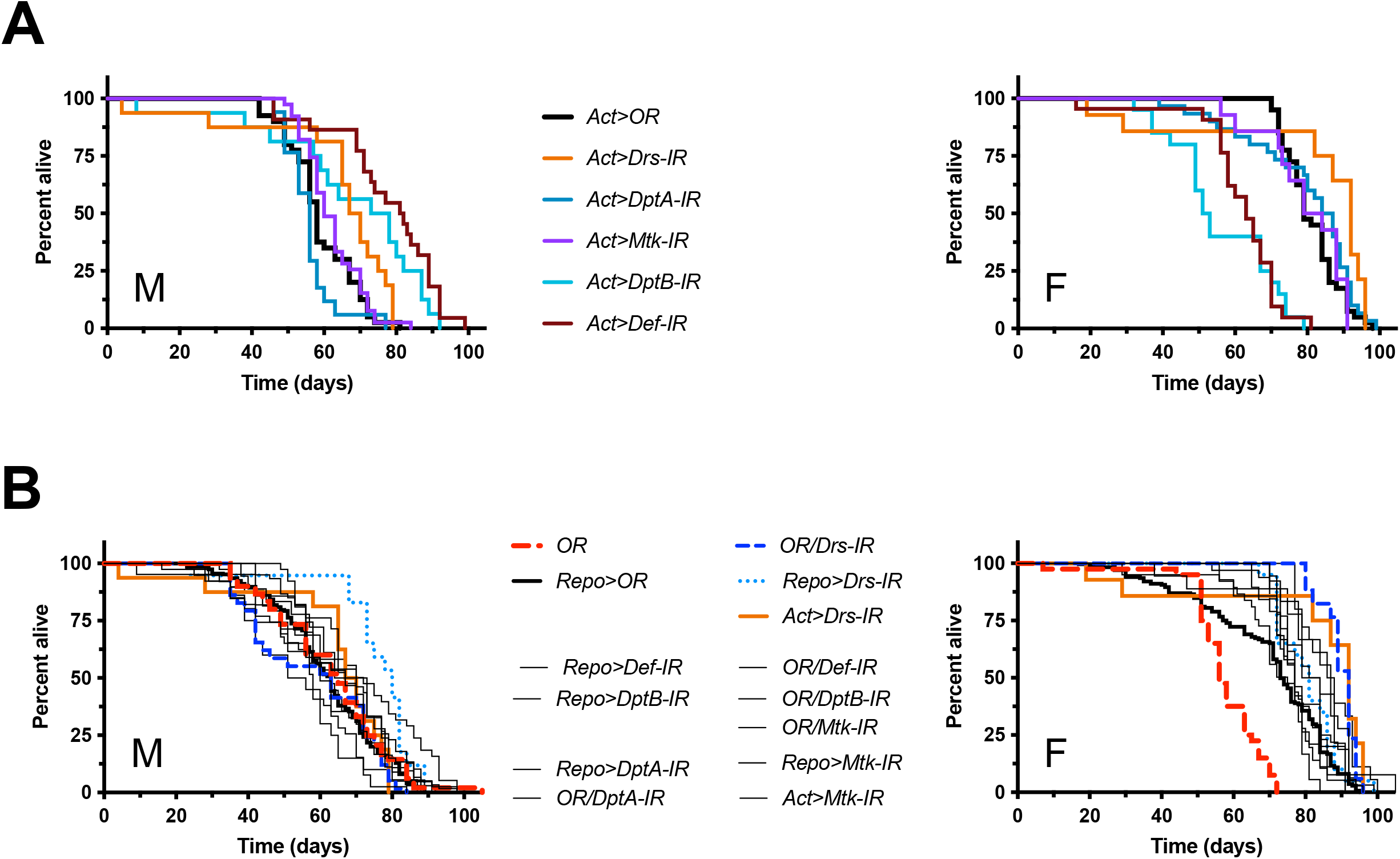
AMP gene silencing ubiquitously or in glia emphasizes the importance of reference genotypes for lifespan context. Males (M) and females (F) shown in separate panels. A) Comparison of multiple *Actin5C-Gal4>AMP-IR* lines shows possible major lifespan effects of Act>Def-IR, with males having extended lifespan compared to controls, while females had reduced lifespan. However these data are exactly opposite to trends from *Def*^*SK3*^ mutant data, where males had reduced lifespan (Fig. 2) and females had comparable lifespan to wild-type (Fig. S3). B) In general, lifespan differences are not especially striking in the context of additional genetic background controls and references. For instance, female *Act>Drs-IR* lifespan is not very different from *OR/Drs-IR* control, suggesting the *Drs-IR* genetic background is long-lived independent of *Drs* knockdown. Similarly, despite a seeming lifespan extension effect by *Act>DptB-IR* in males, we found no lifespan effect in *Dpt*^*SK1*^ or *DptB* mutants, both deficient for *DptB* (Fig. 2).

**Figure S5:**
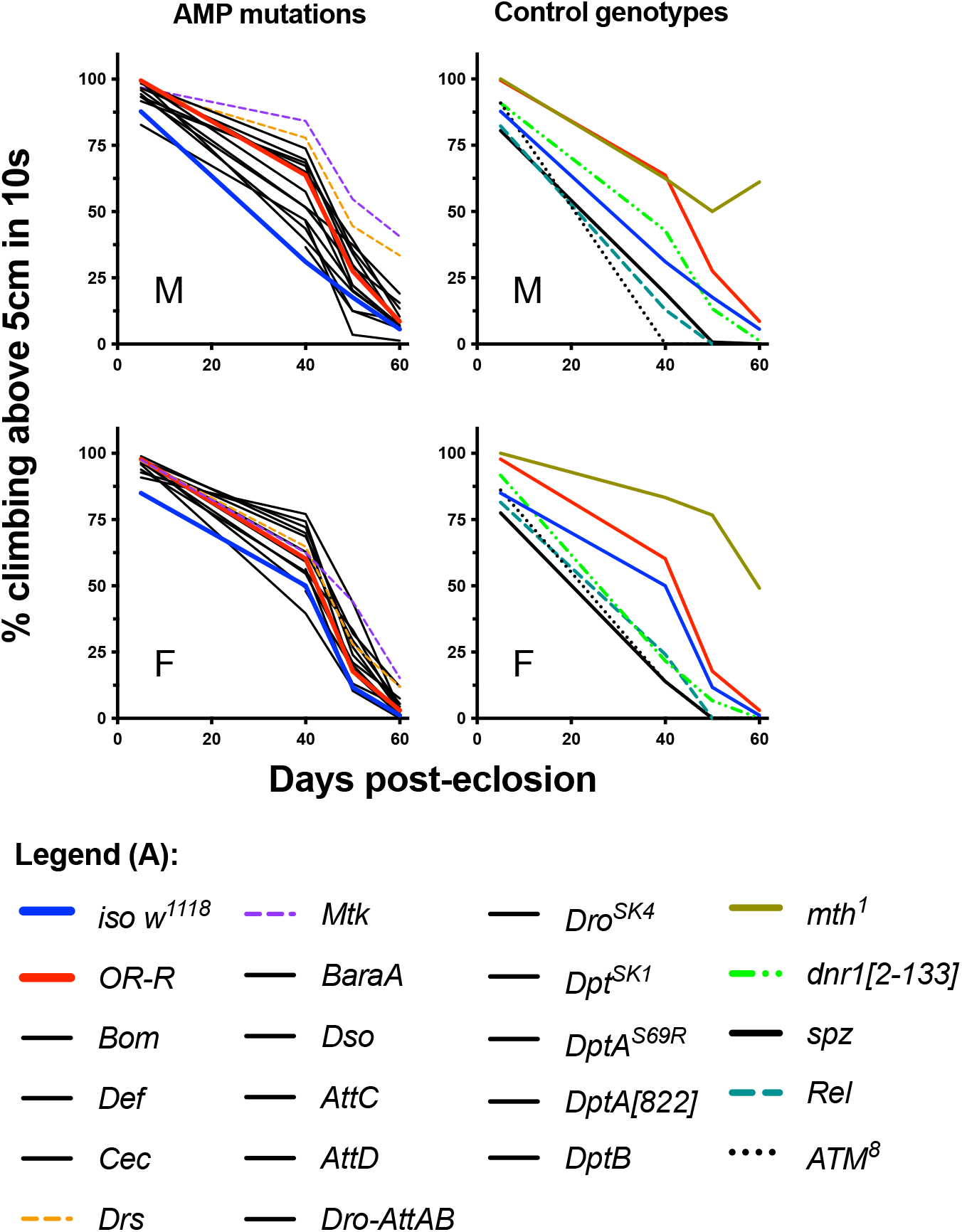
Climbing pass rate data overview from individual mutants. Males (M) and females (F) shown in separate panels. A) Climbing pass rate curves over aging in males and females. AMP mutants displaying climbing rate within wild-type range are left with solid black lines (left panels). Of note, *Drs*^*R1*^ and *Mtk*^*R1*^ are the only mutations which are caused by the insertion of *white*^*+*^ transgenes. Thus, we cannot exclude that their improved climbing effect results from the presence of *white*^*+*^ compared to *iso w*^*1118*^ and other AMP mutants. In line with this idea, *OR-R white*^*+/+*^ wild-type flies similarly trended towards better climbing competence than *iso w*^*1118*^ flies, although many factors likely distinguish these two genetic backgrounds.

**Figure S6:**
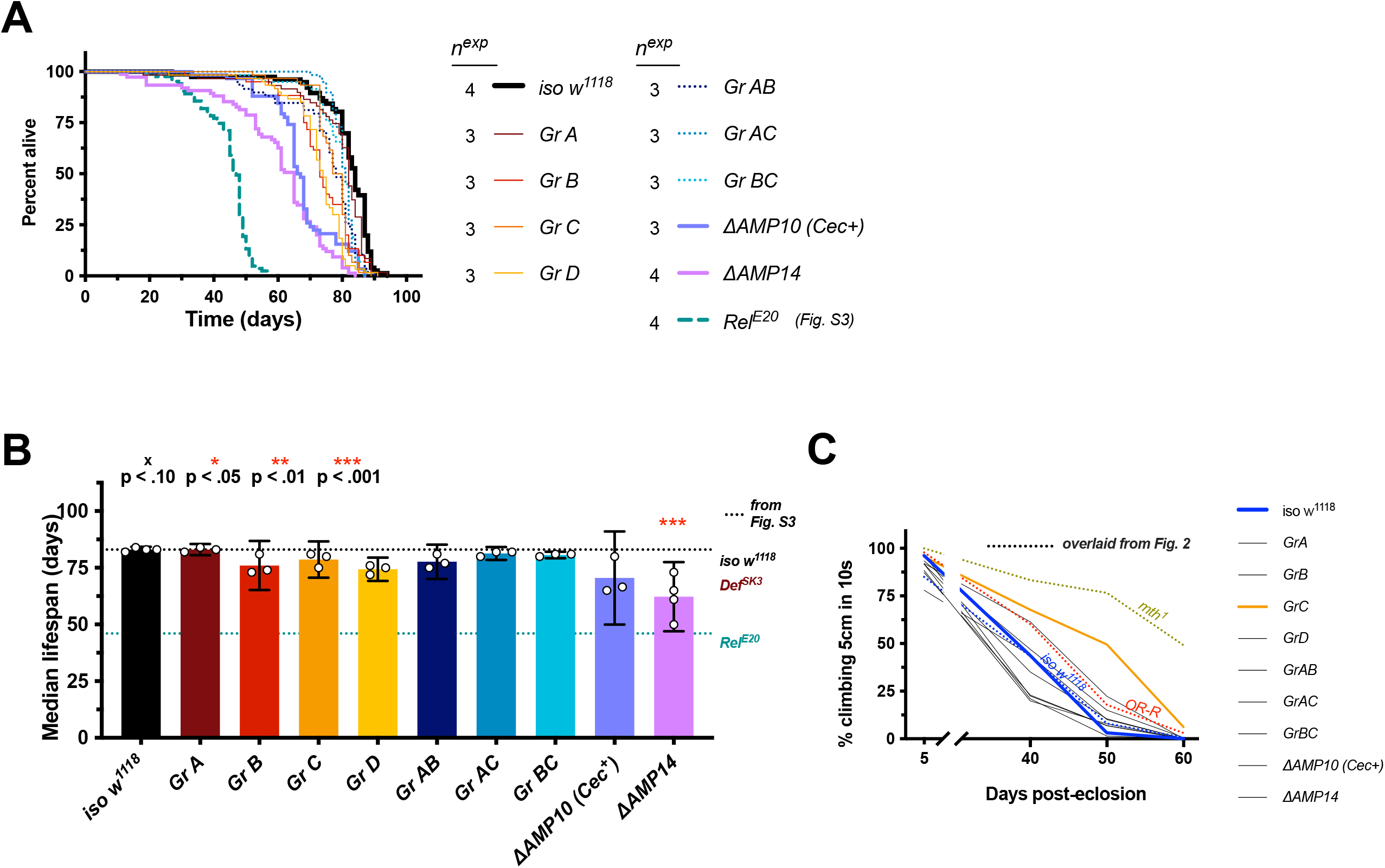
*ΔAMP14* flies have significantly reduced lifespan. Male flies are shown in Figure 3. A) survival curves of various compound AMP mutants. The lifespan of *Rel*^*E20*^ from Fig. S3 is overlaid for direct comparison. B) Median lifespans of compound AMP mutants. Dotted lines indicate average median lifespans from Fig. S3 of *iso w*^*1118*^ (top), *Def*^*SK3*^ alone (middle), and *Rel*^*E20*^ (bottom) for easier comparisons across figures. Statistic summaries (p-value: ^**x**^, *, **, ***) reflect comparisons to *iso w*^*1118*^ data specific to Figure S6. C) Climbing pass rates of AMP group mutants, with climbing curves from genotypes in Fig. S3 overlaid for direction comparison. *Group C* is highlighted for having a slightly improved climbing over aging, though this improvement is still minor compared to the climbing competence of *mth*^*1*^ flies (but see Fig. S5 caption and File S4).

**Figure S7:**
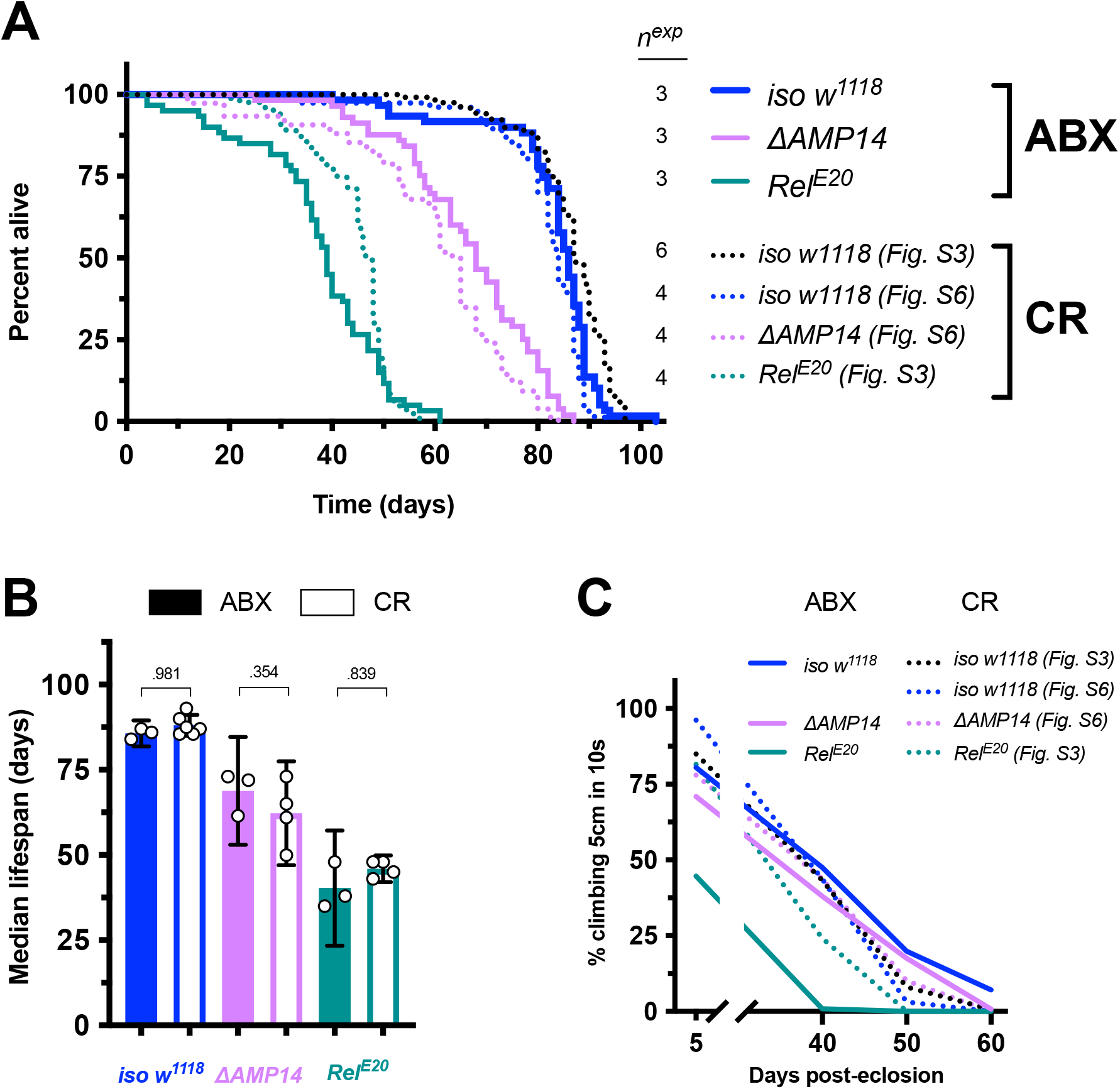
Microbiome depletion rescues the *ΔAMP14* fly lifespan. Female flies are reported here and male flies in Figure 4. A) Survival curves, including both antibiotic-reared flies (ABX), and also conventionally-reared (CR) lifespans from previous figures as dotted lines for direct comparison. B) Median lifespans, including both ABX and CR fly lifespans for direct comparison. (conventionally-reared *iso w*^*1118*^ and *Rel*^*E20*^ lifespans shown in Fig. 2C). C) Climbing pass rates of ABX (solid lines) and CR (dotted lines) flies at 5, 40, 50, and 60 days post-eclosion. Of note, antibiotic treatment did not rescue female Δ*AMP14* lifespan to the same extent as in males (Δ*AMP14-ABX vs*. Δ*AMP14-CR*, P = .354). The general trend of the antibiotic treatment remains the same as in males: Δ*AMP14* flies lived longer in ABX conditions than CR flies (consistent with visual inspection in Figure S7A). Indeed, using a more standard CoxPH mixed model, the rescue effect of antibiotics in Δ*AMP14* females is significant (*P* = .004). Dissecting the effect at the level of sex*genotype interactions in our study should still be interpreted with caution, as any sex*genotype interaction is indistinguishable from ‘vial effects’ due to our experimental design, particularly important as microbiome development in conventionally-reared AMP mutants in more stochastic than control flies (Marra et al., 2021). A full discussion is provided in File S4 (supplemental text).

**Figure S8:**
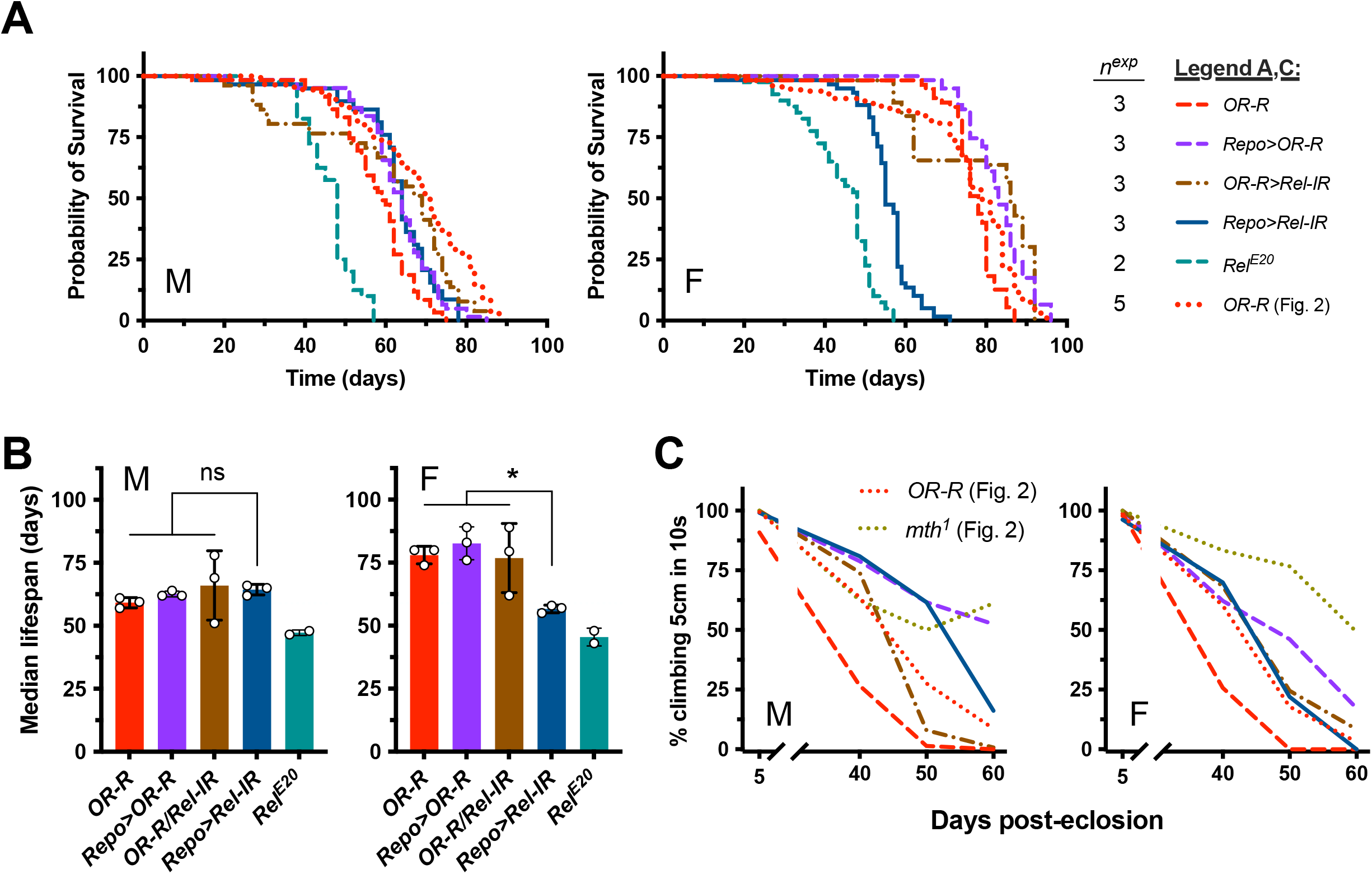
Silencing *Relish* in glia does not extend lifespan. Males (M) and females (F) shown in separate panels. A) Silencing Relish in glia provides no lifespan benefit in males, and was even associated with reduced longevity in females (*Repo>Rel-IR* genotype: *UAS-Rel-IR/+; Repo-Gal4/+*). B) Median lifespans of *Repo* and *Rel-IR* combinations. P < .05 = “*”. C) Climbing pass rates of flies with various combinations of *Repo* and *Rel-IR* systematic crosses showing linear progression of climbing competence loss. The *Repo-Gal4* genetic background seems to improve climbing competence into old age independent of the RNAi construct in males, and perhaps slightly in females. Of note, the lifespan and/or climbing of *OR-R* wild-type flies in these experiments was poor compared to the previous *OR-R* experiments shown in Fig. 2. We therefore included the Fig. 2 *OR-R* lifespan and climbing (and *mth*^*1*^ climbing) in the present figure (A, C) for context, and for ease of direct comparisons.

**Table S1:**
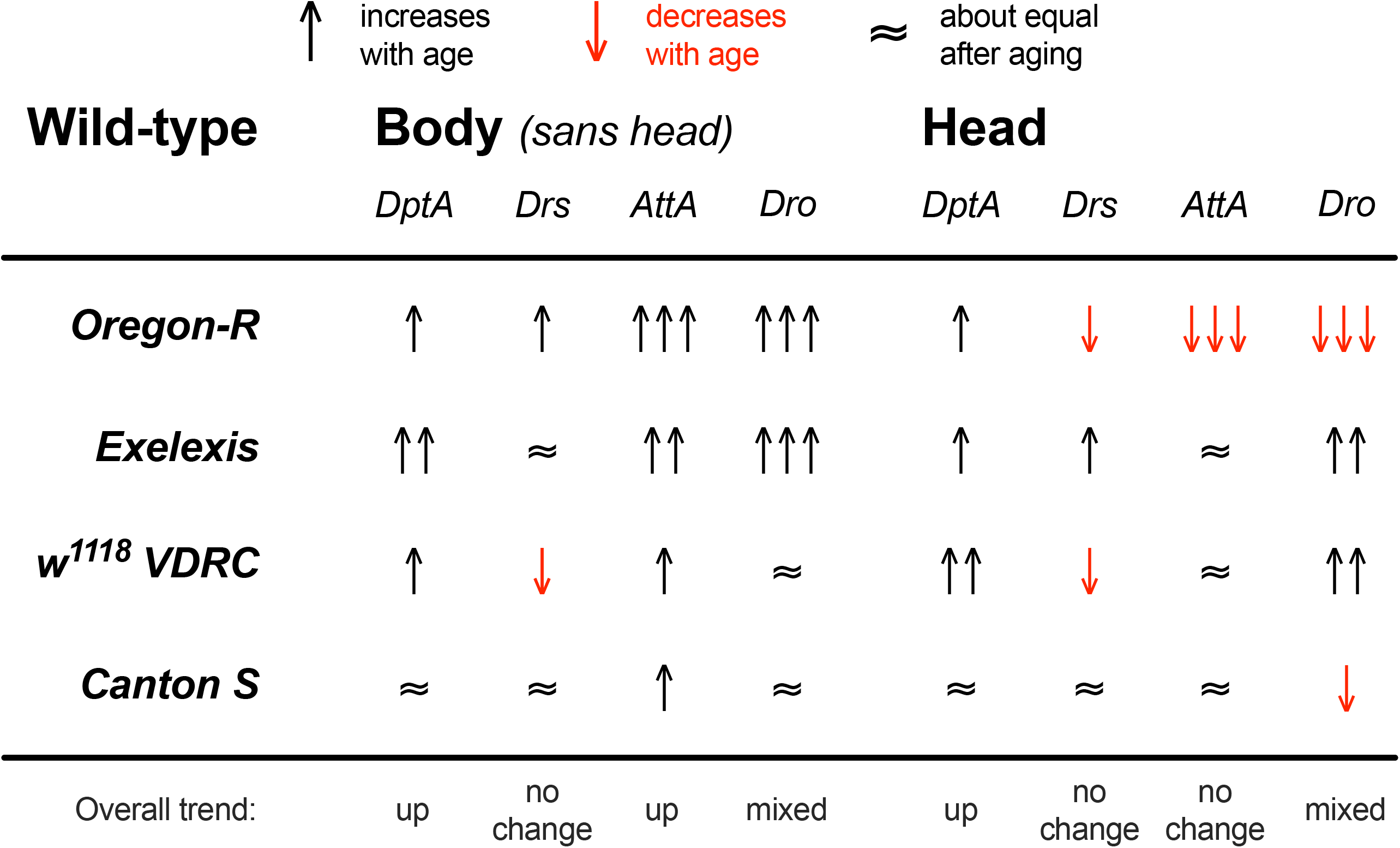
No consistent signal of high AMP expression in the head upon aging. We aged four wild-types (*Oregon-R, Exelexis, w*^*1118*^ *VDRC, and Canton S*) for 40 days and compared AMP expression in 40 day old flies to 5 day old flies, specifically in heads or decapitated bodies. While Imd-responsive AMPs were somewhat consistently upregulated in the body with aging, there was little consistent upregulation of AMPs in the head with aging. When present, upregulation was often minor compared to the expression seen during a systemic immune response for those AMP genes (e.g. at 40dpe *DptA* was induced anywhere from 2-50x in the head, or 5-130x in the body, while induction upon infection often reaches 500-1000x using the same normalization procedure). Underlying data and additional qPCR data from a separate experiment following AMP expression in the heads of aging flies at 8, 15, and 30 days are provided in File S2.

**File S1: Median lifespan and other summary statistics from all genotypes across all experiments, separated by genotype * experiment * sex**. Tabs within this excel file are named according to which experiment they relate to. Full survival raw data are uploaded in File S5.

**File S2: AMP expression data in the head. Contains data related to Table S1 for four wild-type flies collected at 5dpe and 40dpe**. Includes additional data on AMP expression in the heads of aged flies at 8, 15, and 30 days post-eclosion.

**File S3: Description of fly stocks and qPCR primers used in this study**.

**File S4: Supplemental text**. This supplementary discussion addresses issues and limitation of our study that we encountered when combining mutations in the DrosDel isogenic background. We discuss the use of median lifespan as our primary readout. There is also discussion of the unique, but minor, climbing effect of *Mtk* and *Drs* mutation in the light of a recent study, but also with the caveat of *Mtk*^*R1*^ and *Drs*^*R1*^ having a *white*^*+*^ transgene insertion that may affect aging and neurodegeneration. Also contains discussion of the consideration of sex*genotype interactions as vial effects, and how future studies may better assess AMP interactions with sex and aging.

**File S5: Raw data for all experiments**.

## Notes

### Competing Interest Statement

The authors have declared no competing interest.

### Summary of Updates

Revisions have added additional experimental data, and separated the sex-specific lifespans. The manuscript's main figures now show male lifespans, while female lifespans are shown in the supplementary figures. Broadly, trends were similar between males and females in terms of mutation effect.

