## Supplementary material for "Antimicrobial peptides do not directly contribute to aging in *Drosophila*, but improve lifespan by preventing dysbiosis": File S4

***Supplemental text***

*Statistical analysis: CoxPH vs. one-way ANOVA*

Longevity data can be highly variable, as emphasized by median lifespan variance seen in figures 2-4. Of note, dysbiosis onset was also variable in AMP mutants, which may explain some of the high variability we observed. This microbiome variability was further compounded by vial effects, which in our experiments within a genotype are akin to *sex*experimental_replicate* effects as sexes were kept in separate vials. As a result, a single dysbiosis event in one vial in one experiment can show up in our CoxPH model as if it is a sex-specific effect, unique to a certain genotype, most strongly seen in a specific experiment (output as a *sex*experiment*genotype* interaction). The overall significance signal at the genotype level was often swayed by such experiment-specific vial effects, as if a vial effect was evidence of a genotype*sex interaction. Visual inspection of curves across experiments confirmed these significance signals were likely best explained by the high stochasticity of lifespan experiments alongside vial effects, not a genuine *genotype*sex* interaction.

We therefore chose a simple readout of median lifespan per genotype per experiment. This allowed us to present our data in a more easily digested format, and de-emphasizes outlier experiments, focusing on the repeatability of the median lifespan of flies across experiments. A future study, focusing on fewer genotypes and greater total replication within genotypes, will be more appropriate to tease out sex-specific effects of AMP mutations.

*Genetic considerations*

Isogenization is a probabilistic event. As such, while we recombined mutations into the DrosDel isogenic background over seven generations, there is likely variation around how truly “isogenic” these individual AMP mutant flies are. This variance effect is compounded by further recombining mutations in our AMP group approach, which retains the original background locus neighbouring the mutation. Accordingly, our multiple AMP mutation genotypes should have compounding and further increased stochasticity in their isogenic status.

This potential issue is most clear for the *Def^SK3^* lifespan effect. For *Def^SK3^* flies, we observed an allele in the *Drosocin* locus in the original *iso Def^SK3^* flies that did not match the DrosDel isogenic background, encoding the alternate version of a Threonine/Alanine polymorphism in the *Drosocin* gene’s C-terminus peptide Buletin, which mediates defence against *Providencia burhodogranariea* bacteria (Hanson et al., 2022; R Soc B). The distance between *Defensin* and *Drosocin* on chromosome 2R is ~4.7 million base pairs, which suggests a low level of isogenization in the original *iso Def^SK3^* line used in Parvy et al. (2019; eLife) and Hanson et al. (2019; eLife). We performed additional recombination of *Def^SK3^* with the DrosDel background, which did not change individual mutant lifespan. However, the extra-recombination *Def^SK3^* chromosome, in combination with *ΔCec^A-C^* to generate AMP *Group A* flies, showed markedly improved lifespan compared to the original *Def^SK3^* chromosome. Median lifespan differences were: original *Group A* (62.8 days) vs. additionally recombined *Def^SK3^ Group A* (74.3 days). This striking lifespan improvement from additionally recombining the *Def^SK3^* mutation into the DrosDel background, alongside the lack of impact of *Defensin* RNAi, further emphasizes a cautious interpretation of the *Def^SK3^* effect (see File S1 “Fig3supp_GrA1_vs_GrA2”).

*Effect of Mtk^R1^ and Drs^R1^ on climbing competence*

We observed slightly improved climbing competence of *Mtk* and *Drs* mutants individually (Fig. 2D and Fig. S3) and *Mtk, Drs* double mutants (Fig. 3D). A previous study also suggested that the same *Mtk* and *Drs* stocks used here have countervailing effects on outcome of traumatic brain injury when combined as AMP *Group C* (Swanson et al., 2020; G3). In general, we do not focus on these differences, as they were minor compared to *methuselah* fly strain, which set an upper bound for possible climbing effects. However these results taken together present an interesting hypothesis that *Mtk* and *Drs* impact brain health, including over aging.

However, *Mtk^R1^* and *Drs^R1^* are the only AMP mutations generated by our group by homologous recombination (Baena-Lopez et al., 2013; Development), involving a *white^+^* rescue construct. Not only does the *white* gene regulate gut dysplasia with aging (Sasaki et al., 2021; Nat Metab), but the *w^1118^* mutation may also cause retinal neurodegeneration and reduced climbing compared to *white^+^* controls (Ferreiro et al., 2018; Front Neurosci). Here, *methuselah* mutants (Fig. 2), alongside *ΔAMP14* flies reared conventionally or with antibiotics that have different lifespan but similar climbing (Fig. 4), demonstrate that lifespan and climbing competence can be independent phenotypes. Given a possible positive climbing competence/aging effect of the *Repo-Gal4* background alone (Fig. S4), future experiments disrupting AMPs or Imd signalling in the brain should be interpreted carefully.

*Considering sex in interactions of AMPs and aging in our study*

AMPs assuredly regulate the microbiome, as demonstrated in a number of studies across organisms (Bosch and Zasloff, 2021), including our own using *∆AMP14* flies (Marra et al., 2021). Indeed one of our major findings from Marra et al. (Marra et al., 2021) was that community structures changed in AMP mutants, and there was increased variance in the microbiome. The bottlenecking of 20 individuals into a fly community that then develops over 3 months of aging means that small differences in initial vial conditions can snowball into major differences across vials over the course of a lifespan experiment: so-called “vial effects.” Indeed one of the clear signals in our median lifespan data is that AMP mutants have increased variance in median lifespan compared to *iso w^1118^*.

We kept sexes separate over the course of the aging experiment. As a result, within an experiment the denomination of male or female is also associated with being assigned to separate vials, subject to their own vial effects. We are therefore cautious of over-interpreting our data at the sex*genotype level, as there is a clear signal of increased stochasticity in AMP mutant lifespans in general, and so it is more likely that vial effects will randomly generate samples at both ends of the potential spectrum of microbiome diversity and abundance. With a small sampling of only two vials (one male, one female) per genotype across three replicate experiments in most cases, it is too preliminary to say if AMPs have sex-specific interactions with aging. Of note, our *Def^SK3^* mutation caused a more male-specific lifespan reduction (Fig. 2A vs. Fig. XX), but then initial *Defensin* RNAi experiments gave the exact opposite result, with extended male lifespan and reduced female lifespan (Fig. S2), which, even granting genetic background effects, suggests concluding a positive or negative impact of *Defensin* on lifespan overall, or in sex-specific fashion, is unlikely to be true in a general sense. Indeed, by screening many genotypes, we not only perform many hypothesis tests, but we are specifically screening genotypes expected to have a higher stochasticity in lifespan by virtue of their stochastic microbiome development (Marra et al., 2021). This may make our analyses more prone to false positive significance tests when comparing genotypes, compared to aging studies that do not disrupt key immune genes for regulating the microbiome. A future study on sex by AMP interactions would benefit from a male-female co-housing design to homogenize the environments of males and females, ensuring that vial effects do not confound sex*genotype interactions.
