## Supplementary material for "Antimicrobial peptides do not directly contribute to aging in *Drosophila*, but improve lifespan by preventing dysbiosis": File S5: Fig1A.R

```
### Useful link http://anson.ucdavis.edu/~hiwang/teaching/10fall/R_tutorial%201.pdf

setwd("~/Desktop/Lemaitre Lab/Aging paper OFFLINE/Revisions/Figures_SuppFiles/FileS5_rawdata/Fig1/Fig1A")

library(survival) # R only
#install.packages("colorspace")
library(colorspace)
#pal<-choose_palette()
library(hues)

library(readxl)
#FlyData <- read_excel("/Volumes/Lemaitre-Lab/Mark Hanson/Mark-Infections/2020/Longevity/Baramycin CoxPH/Baramycin-CoxPH_mf.xls")
FlyData <- read_excel("~/Desktop/Lemaitre Lab/Aging paper OFFLINE/Revisions/Figures_SuppFiles/FileS5_rawdata/Fig1/Fig1A/Fig1A.xls")
#FlyData <- read_excel("/Volumes/Lemaitre-Lab/Mark Hanson/Mark-Infections/2020/Longevity/Baramycin CoxPH/Baramycin-CoxPH_males.xlsx")
#FlyData <- read.csv(file.choose(), header=T)
head(FlyData)

#########################################################
###Adding to get median lifespans ***PER EXPERIMENT***###
#########################################################
#GtypeExp<-paste(FlyData$Genotype,FlyData$Experiment,FlyData$Sex)#,FlyData$Sex)
#FlyData$Genotype<-GtypeExp

Gtype<-as.factor(FlyData$Genotype) #Note: it will organize levels alphabetically first by capitals then by lower case
levels(Gtype)
Gtype2<-unique(c(as.character(levels(Gtype)))) #Sorts genotypes by alphabetical order, doesn't require you to enter them in alphabetical order!
Life<-FlyData$Time2 #Life where time can't be -40000ish
Status<-FlyData$Censored #Censoring status, 0=censored
Sex<-FlyData$Sex
Exp<-FlyData$Experiment

###################################################
#Convert R survival data to Graphpad format
#Answer provided by Sotos in https://stackoverflow.com/questions/47770488/like-dcast-but-without-sum-of-data

###Prepare dataframe for export###
FlyData$Time.dead <- NULL #Get rid of extraneous column
Gtype<-as.factor(FlyData$Genotype); Life<-FlyData$Time2 ; Status<-as.numeric(FlyData$Censored)
FlyData<-data.frame(Gtype,Life,Status)

###Convert to Graphpad format###
#install.packages("backports")
library(backports)
library(tidyverse)
df<-FlyData %>% mutate(new = row_number()) %>% spread(Gtype, Status, fill = NA) %>% select(-new)
write.csv(df, file="Fig1A_earlysurvs.csv")
###################################################

### Kaplan-Meier test statistics and survival plot separated by Treatment
quartz(h=6,w=9) ## The equivalent for a PC is windows()
par(mfrow=c(1,1),mar=par()$mar+c(0,0,0,7), pty="s", xpd=TRUE)
flysurv <- survfit( Surv(Life, Status)~ strata(Gtype))
plot(flysurv, lty=c(1:3), lwd=c(1.5), col=iwanthue(20), xlab="Time (days)", ylab="Survival Probability", mark=c(1:20), xaxt="n", yaxt="n")#conf.times=c(1,2,3,4,5,6,7), conf.cap=.005
points(flysurv,col=iwanthue(20),pch=c(1:20))
axis(side=1,at=seq(0,100,by=10), cex.axis=0.8)
axis(side=2,at=seq(0,1,by=.25))
legend(110,1, (as.factor(Gtype2)), lty=c(1:3),lwd=c(1.5), col=iwanthue(20), cex = 0.8, pch=c(1:20)) #set legend to x=16 for graphs with xlim 14, and 8 for graphs with xlim 7
title(main="25'C early experiments",cex.main=0.8,line=1)
text(165,0.5,"coxph(formula = Surv(Life, Status) ~ Gtype + Exp + Sex, method = efron)
AMPChr2 as baseline,  n= 479, number of events= 398 

                  coef exp(coef) se(coef)      z Pr(>|z|)    
Gtypeiso w1118  2.4095   11.1285   0.1553 15.515  < 2e-16 ***
GtypeOR-R       0.6803    1.9745   0.1355  5.022 5.11e-07 ***
Exp16-07-2018   0.5535    1.7394   0.1543  3.587 0.000334 ***
Exp21-09-2017   1.4733    4.3635   0.1359 10.843  < 2e-16 ***
Sexm            0.3751    1.4551   0.1058  3.545 0.000392 ***Exp ***, Sex ***",cex=0.4,pos=2) #set x to 25 for graphs with xlim = c(0,14), and 12.5 for graphs with xlim = c(0,7)

full.coxph <- coxph(Surv(Life, Status)~Gtype+Exp +Sex,method='efron')
summary(full.coxph)
#Exp significant, and with Exp in model AMPs not significant
flysurv


survoutput<-summary(flysurv)$table
write.csv(survoutput,file="survoutput.csv")
```
