## Supplementary material for "Antimicrobial peptides do not directly contribute to aging in *Drosophila*, but improve lifespan by preventing dysbiosis": File S5: nora_revisions.pdf

**Longevity, 25'C, individual iso flies by experiment**
**2017-2021 experiments (7 in total, updated Oct 27th 2021)**

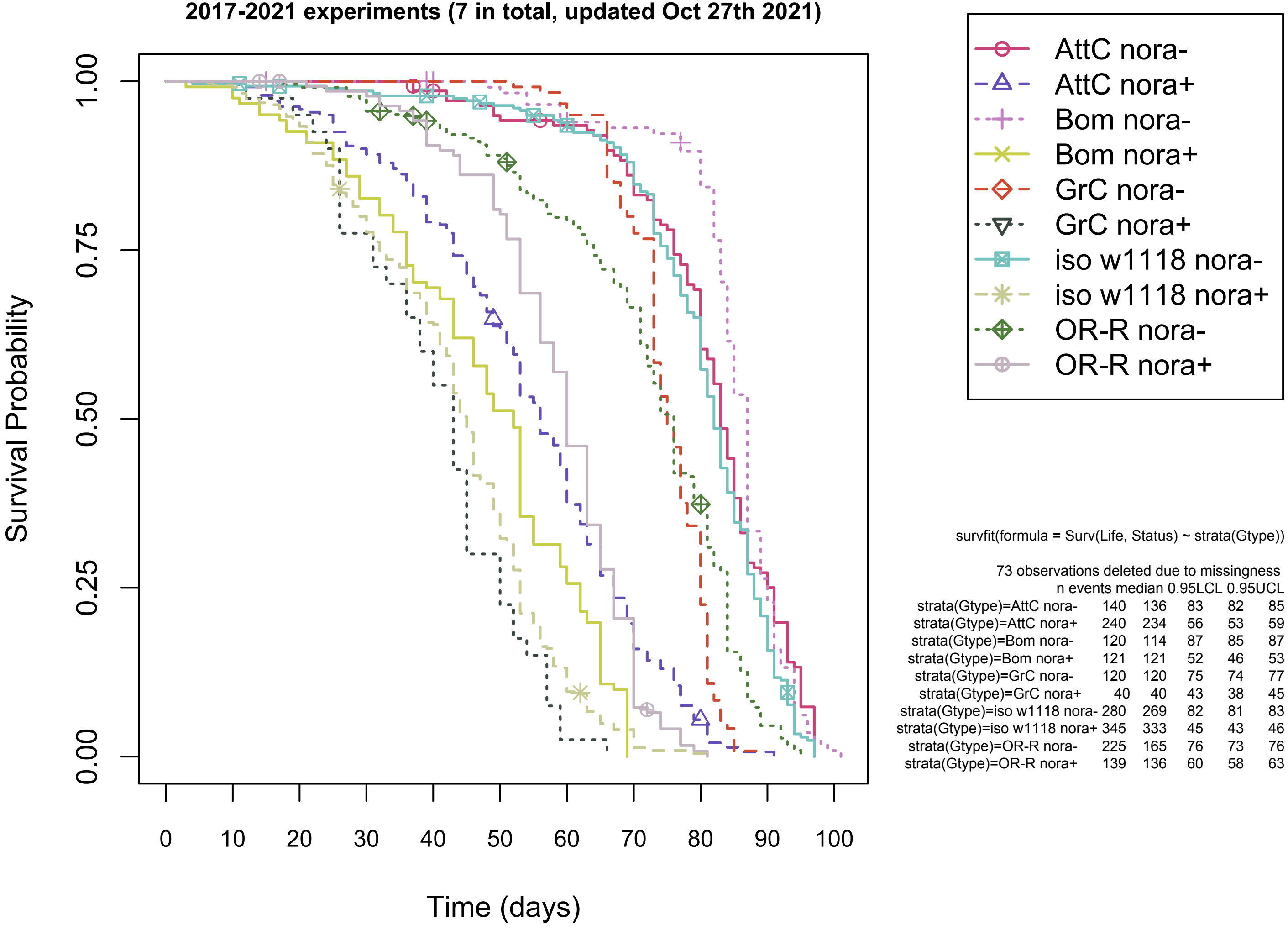


survfit(formula = Surv(Life, Status) ~ strata(Gtype))

73 observations deleted due to missingness

| | n | events | median | 0.95LCL | 0.95UCL |
|---|---|---|---|---|---|
| strata(Gtype)=AttC nora- | 140 | 136 | 83 | 82 | 85 |
| strata(Gtype)=AttC nora+ | 240 | 234 | 56 | 53 | 59 |
| strata(Gtype)=Bom nora- | 120 | 114 | 87 | 85 | 87 |
| strata(Gtype)=Bom nora+ | 121 | 121 | 52 | 46 | 53 |
| strata(Gtype)=GrC nora- | 120 | 120 | 75 | 74 | 77 |
| strata(Gtype)=GrC nora+ | 40 | 40 | 43 | 38 | 45 |
| strata(Gtype)=iso w1118 nora- | 280 | 269 | 82 | 81 | 83 |
| strata(Gtype)=iso w1118 nora+ | 345 | 333 | 45 | 43 | 46 |
| strata(Gtype)=OR-R nora- | 225 | 165 | 76 | 73 | 76 |
| strata(Gtype)=OR-R nora+ | 139 | 136 | 60 | 58 | 63 |
