## Supplementary material for "Antimicrobial peptides do not directly contribute to aging in *Drosophila*, but improve lifespan by preventing dysbiosis": File S5: ind mf CoxPH.pdf

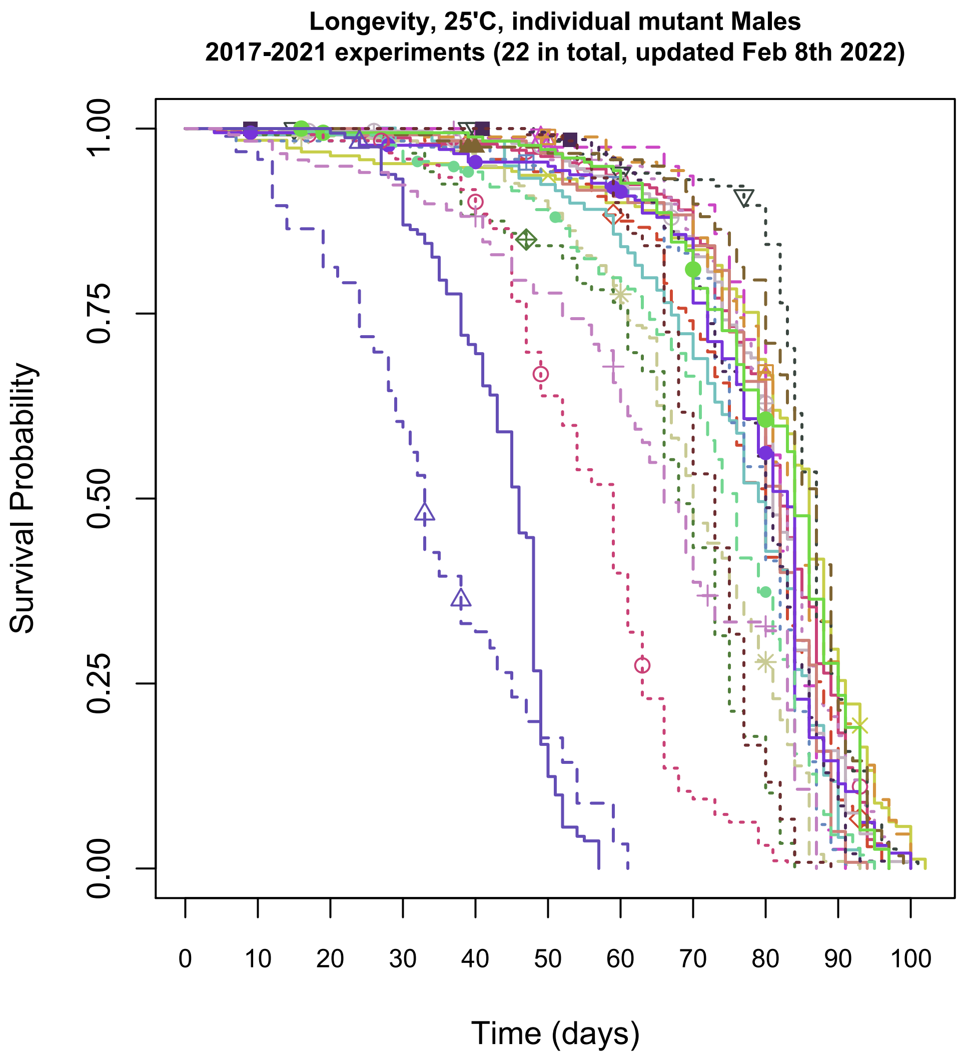


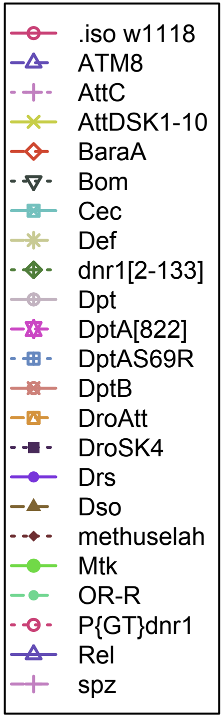


coxph(formula = Surv(Life, Status) ~ Gtype + Exp + Sex, method = efron)
iso as baseline,n= 3650, number of events= 3265

| | coef | exp(coef) | se(coef) | z | Pr(>\|z\|) | |
|---|---|---|---|---|---|---|
| GtypeATM8 | 4.36847 | 78.92286 | 0.17400 | 25.106 | < 2e-16 | *** |
| GtypeAttC | 0.07876 | 1.08194 | 0.11820 | 0.666 | 0.50520 | |
| GtypeAttDSK1-10 | -0.45647 | 0.63352 | 0.10808 | -4.224 | 2.41e-05 | *** |
| GtypeBaraA | 0.08853 | 1.09257 | 0.11683 | 0.758 | 0.44857 | |
| GtypeBom | -0.58000 | 0.55990 | 0.12121 | -4.785 | 1.71e-06 | *** |
| GtypeCec | 0.60398 | 1.82938 | 0.11950 | 5.054 | 4.32e-07 | *** |
| GtypeDef | 1.30132 | 3.67414 | 0.12198 | 10.668 | < 2e-16 | *** |
| Gtypednr1[2-133] | 1.35426 | 3.87390 | 0.12665 | 10.693 | < 2e-16 | *** |
| GtypeDpt | 0.29893 | 1.34842 | 0.10866 | 2.751 | 0.00594 | ** |
| GtypeDptA[822] | 0.20664 | 1.22954 | 0.14776 | 1.398 | 0.16197 | |
| GtypeDptAS69R | 0.38101 | 1.46376 | 0.12436 | 3.064 | 0.00218 | ** |
| GtypeDptB | 0.07440 | 1.07724 | 0.12416 | 0.599 | 0.54901 | |
| GtypeDroAtt | 0.12400 | 1.13202 | 0.12397 | 1.000 | 0.31719 | |
| GtypeDroSK4 | 0.30686 | 1.35914 | 0.10329 | 2.971 | 0.00297 | ** |
| GtypeDrs | 0.52201 | 1.68541 | 0.12702 | 4.110 | 3.96e-05 | *** |
| GtypeDso | -0.47547 | 0.62159 | 0.11861 | -4.009 | 6.11e-05 | *** |
| Gtypemethuselah | 1.02345 | 2.78278 | 0.12163 | 8.414 | < 2e-16 | *** |
| GtypeMtk | 0.29003 | 1.33647 | 0.12284 | 2.361 | 0.01822 | * |
| GtypeOR-R | 1.16849 | 3.21712 | 0.13266 | 8.808 | < 2e-16 | *** |
| GtypeP{GT}dnr1 | 2.80454 | 16.51954 | 0.14780 | 18.975 | < 2e-16 | *** |
| GtypeRel | 3.20125 | 24.56325 | 0.19978 | 16.024 | < 2e-16 | *** |
| Gtypespz | 1.65301 | 5.22266 | 0.13985 | 11.820 | < 2e-16 | *** |

Exp = *** Sex = ***
