## Supplementary material for "Antimicrobial peptides do not directly contribute to aging in *Drosophila*, but improve lifespan by preventing dysbiosis": File S5: Fig3.pdf

**Longevity, 25'C, compound mutant Males + Females**
**2017-2022 cumulative experiments**

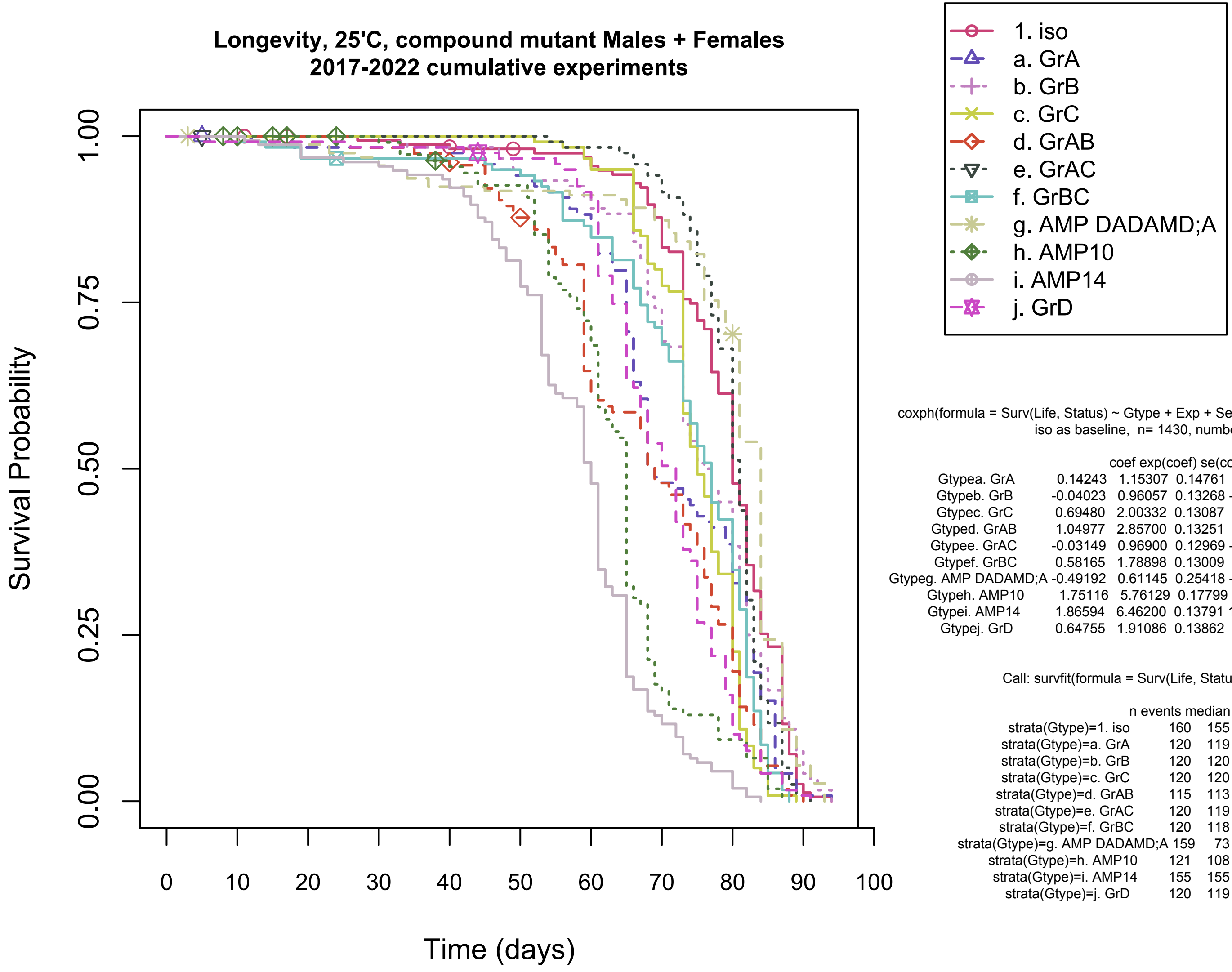


coxph(formula = Surv(Life, Status) ~ Gtype + Exp + Sex, method = efron)
iso as baseline, n= 1430, number of events= 1319

| | coef | exp(coef) | se(coef) | z | Pr(>\|z\|) | |
|---|---|---|---|---|---|---|
| Gtypea. GrA | 0.14243 | 1.15307 | 0.14761 | 0.965 | 0.334586 | |
| Gtypeb. GrB | -0.04023 | 0.96057 | 0.13268 | -0.303 | 0.761713 | |
| Gtypec. GrC | 0.69480 | 2.00332 | 0.13087 | 5.309 | 1.10e-07 | *** |
| Gtyped. GrAB | 1.04977 | 2.85700 | 0.13251 | 7.922 | 2.33e-15 | *** |
| Gtypee. GrAC | -0.03149 | 0.96900 | 0.12969 | -0.243 | 0.808131 | |
| Gtypef. GrBC | 0.58165 | 1.78898 | 0.13009 | 4.471 | 7.78e-06 | *** |
| Gtypeg. AMP DADAMD;A | -0.49192 | 0.61145 | 0.25418 | -1.935 | 0.052949 | . |
| Gtypeh. AMP10 | 1.75116 | 5.76129 | 0.17799 | 9.839 | < 2e-16 | *** |
| Gtypei. AMP14 | 1.86594 | 6.46200 | 0.13791 | 13.530 | < 2e-16 | *** |
| Gtypej. GrD | 0.64755 | 1.91086 | 0.13862 | 4.672 | 2.99e-06 | *** |

Exp***, Sex***

Call: survfit(formula = Surv(Life, Status) ~ strata(Gtype))

| | n | events | median | 0.95LCL | 0.95UCL |
|---|---|---|---|---|---|
| strata(Gtype)=1. iso | 160 | 155 | 80.0 | 80 | 82 |
| strata(Gtype)=a. GrA | 120 | 119 | 69.0 | 68 | 79 |
| strata(Gtype)=b. GrB | 120 | 120 | 76.5 | 73 | 80 |
| strata(Gtype)=c. GrC | 120 | 120 | 75.0 | 74 | 77 |
| strata(Gtype)=d. GrAB | 115 | 113 | 69.0 | 67 | 75 |
| strata(Gtype)=e. GrAC | 120 | 119 | 81.0 | 80 | 82 |
| strata(Gtype)=f. GrBC | 120 | 118 | 77.0 | 74 | 80 |
| strata(Gtype)=g. AMP DADAMD;A | 159 | 73 | 84.0 | 81 | 84 |
| strata(Gtype)=h. AMP10 | 121 | 108 | 65.0 | 62 | 65 |
| strata(Gtype)=i. AMP14 | 155 | 155 | 60.0 | 59 | 61 |
| strata(Gtype)=j. GrD | 120 | 119 | 72.0 | 68 | 73 |
