## Supplementary material for "Antimicrobial peptides do not directly contribute to aging in *Drosophila*, but improve lifespan by preventing dysbiosis": File S5: Fig4.pdf

### Longevity, 25'C, antibiotic-reared Males + Females
### 2021-2022 cumulative experiments

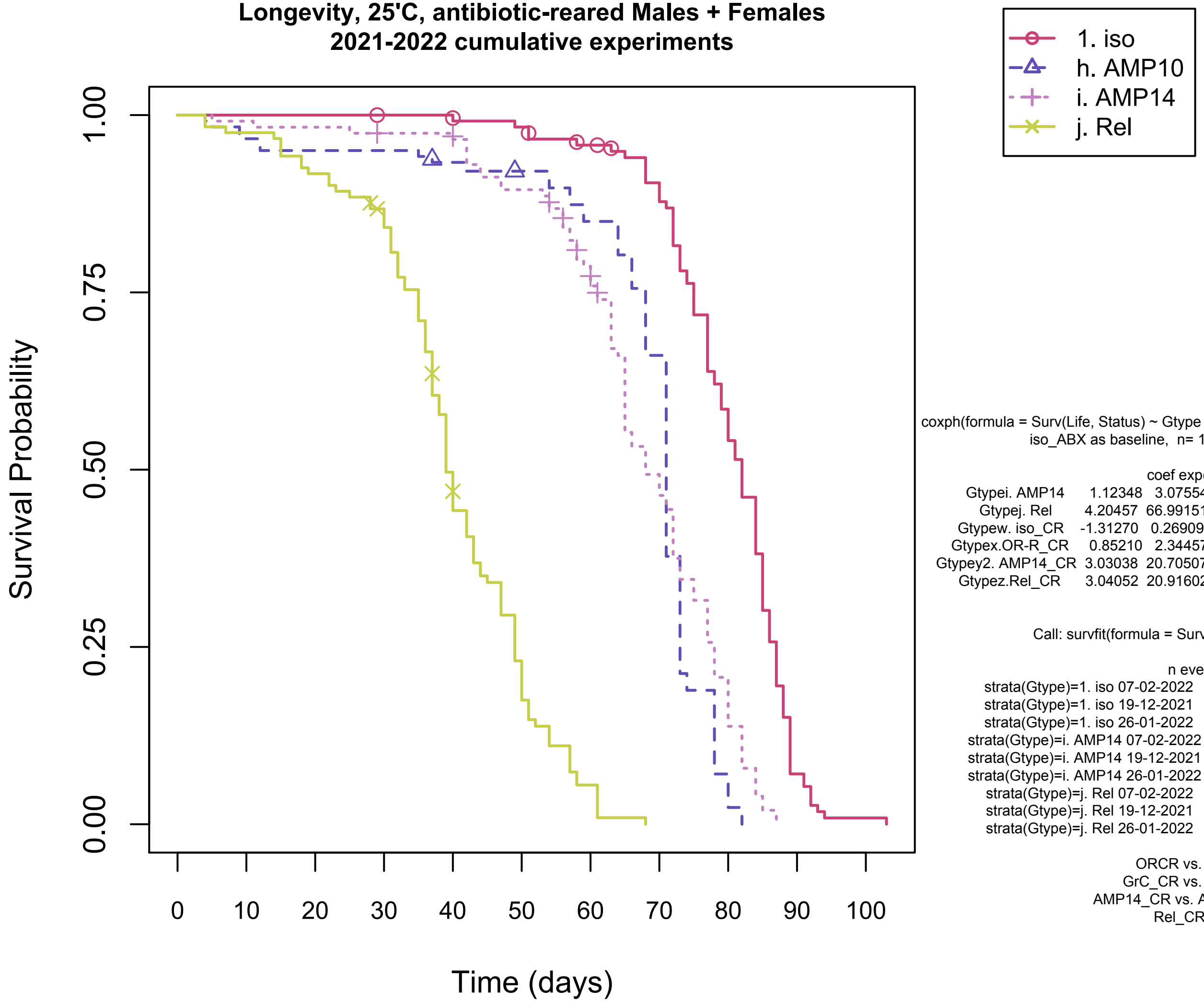


coxph(formula = Surv(Life, Status) ~ Gtype + Exp + Sex, method = efron)
iso_ABX as baseline, n= 1400, number of events= 1296

| | coef | exp(coef) | se(coef) | z | Pr(>\|z\|) | |
|---|---|---|---|---|---|---|
| Gtypei. AMP14 | 1.12348 | 3.07554 | 0.14028 | 8.009 | 1.16e-15 | *** |
| Gtypej. Rel | 4.20457 | 66.99151 | 0.16619 | 25.300 | < 2e-16 | *** |
| Gtypew. iso_CR | -1.31270 | 0.26909 | 0.36103 | -3.636 | 0.000277 | *** |
| Gtypex.OR-R_CR | 0.85210 | 2.34457 | 0.20236 | 4.211 | 2.54e-05 | *** |
| Gtypey2. AMP14_CR | 3.03038 | 20.70507 | 0.22891 | 13.239 | < 2e-16 | *** |
| Gtypez.Rel_CR | 3.04052 | 20.91602 | 0.42249 | 7.197 | 6.17e-13 | *** |

Sex ***, Exp ***

Call: survfit(formula = Surv(Life, Status) ~ strata(Gtype))

| | n | events | median | 0.95LCL | 0.95UCL |
|---|---|---|---|---|---|
| strata(Gtype)=1. iso 07-02-2022 | 40 | 37 | 79.0 | 77 | 84 |
| strata(Gtype)=1. iso 19-12-2021 | 40 | 38 | 85.0 | 82 | 87 |
| strata(Gtype)=1. iso 26-01-2022 | 40 | 38 | 82.0 | 77 | 84 |
| strata(Gtype)=i. AMP14 07-02-2022 | 40 | 37 | 65.0 | 63 | 68 |
| strata(Gtype)=i. AMP14 19-12-2021 | 37 | 31 | 73.0 | 71 | 78 |
| strata(Gtype)=i. AMP14 26-01-2022 | 40 | 36 | 72.0 | 65 | 80 |
| strata(Gtype)=j. Rel 07-02-2022 | 40 | 37 | 35.0 | 32 | 37 |
| strata(Gtype)=j. Rel 19-12-2021 | 40 | 37 | 40.0 | 36 | 47 |
| strata(Gtype)=j. Rel 26-01-2022 | 41 | 38 | 49.0 | 44 | 51 |

ORCR vs. AMP14_ABX, t.test p=0.3893
GrC_CR vs. AMP14_ABX, t.test p=0.0904
AMP14_CR vs. AMP14_ABX, t.test p=0.04574
Rel_CR vs. Rel_ABX, t.test p=0.2368
