## Supplementary material for "Antimicrobial peptides do not directly contribute to aging in *Drosophila*, but improve lifespan by preventing dysbiosis": File S5: Cox_29C_mf.pdf

**Longevity, 29'C, Males + Females**
**2022 three experiments**

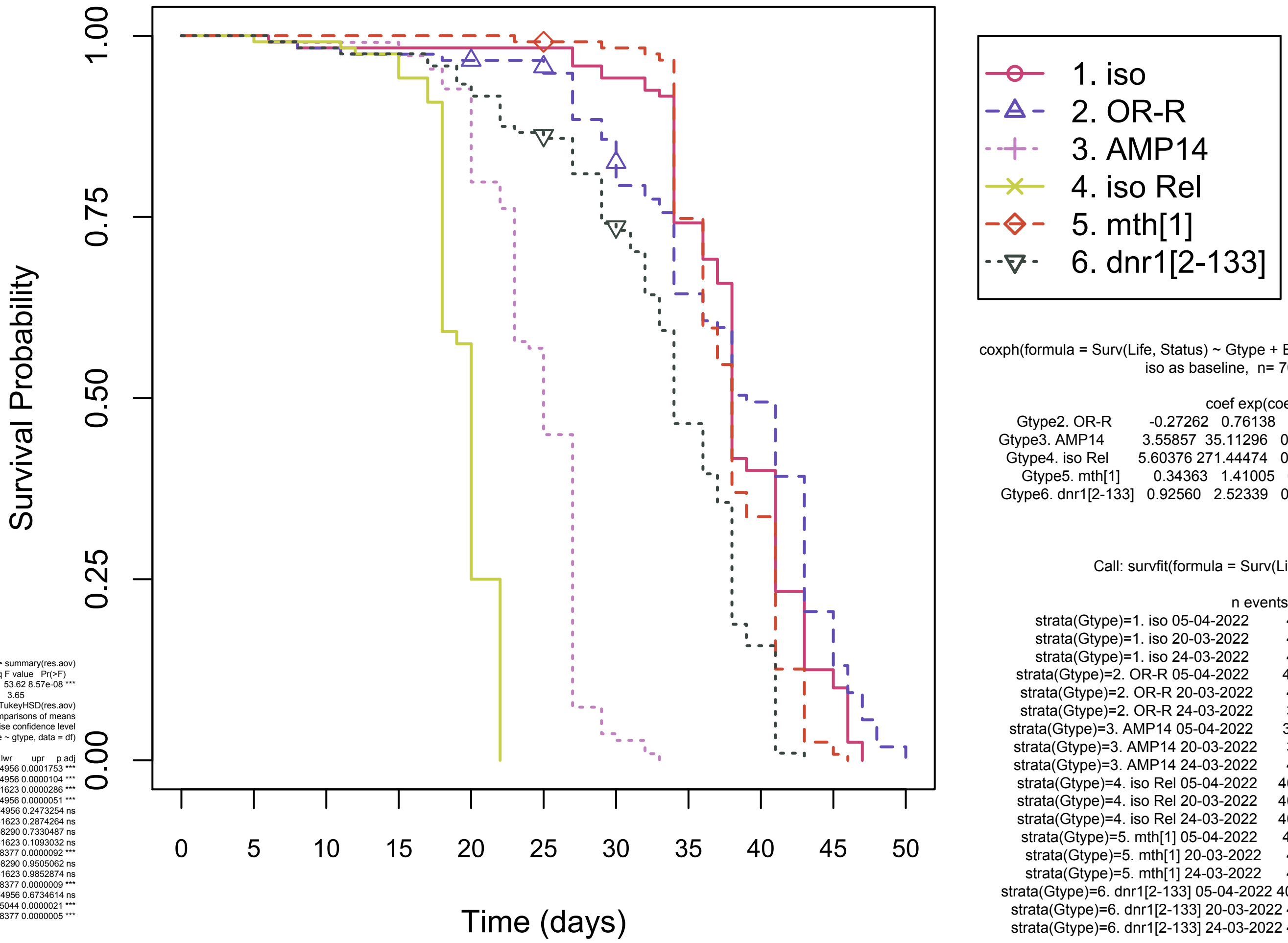


```
coxph(formula = Surv(Life, Status) ~ Gtype + Exp + Sex, method = efron)
iso as baseline,  n= 707, number of events= 680

                        coef exp(coef)  se(coef)      z Pr(>|z|)
Gtype2. OR-R        -0.27262   0.76138   0.13749 -1.983   0.0474 *
Gtype3. AMP14        3.55857  35.11296   0.19711 18.053  < 2e-16 ***
Gtype4. iso Rel      5.60376 271.44474   0.25954 21.591  < 2e-16 ***
Gtype5. mth[1]       0.34363   1.41005   0.13249  2.594   0.0095 **
Gtype6. dnr1[2-133]  0.92560   2.52339   0.13997  6.613 3.77e-11 ***
Exp***, Sex**
```

```
Call: survfit(formula = Surv(Life, Status) ~ strata(Gtype))

                                    n events median 0.95LCL 0.95UCL
strata(Gtype)=1. iso 05-04-2022     40     36   38.0      36      41
strata(Gtype)=1. iso 20-03-2022     40     40   38.0      38      41
strata(Gtype)=1. iso 24-03-2022     40     40   38.0      34      43
strata(Gtype)=2. OR-R 05-04-2022    40     23   41.0      34      NA
strata(Gtype)=2. OR-R 20-03-2022    40     36   38.0      38      43
strata(Gtype)=2. OR-R 24-03-2022    38     34   41.0      34      43
strata(Gtype)=3. AMP14 05-04-2022   30     30   21.0      20      NA
strata(Gtype)=3. AMP14 20-03-2022   39     39   27.0      27      27
strata(Gtype)=3. AMP14 24-03-2022   40     40   23.0      23      25
strata(Gtype)=4. iso Rel 05-04-2022 40     40   20.0      NA      NA
strata(Gtype)=4. iso Rel 20-03-2022 40     40   22.0      NA      NA
strata(Gtype)=4. iso Rel 24-03-2022 40     40   18.0      NA      NA
strata(Gtype)=5. mth[1] 05-04-2022  40     39   38.0      36      NA
strata(Gtype)=5. mth[1] 20-03-2022  40     40   38.0      38      41
strata(Gtype)=5. mth[1] 24-03-2022  40     40   34.0      34      37
strata(Gtype)=6. dnr1[2-133] 05-04-2022 40 24   35.0*     36      NA
strata(Gtype)=6. dnr1[2-133] 20-03-2022 40 40   34.5      31      38
strata(Gtype)=6. dnr1[2-133] 24-03-2022 40 40   34.0      34      37
```

```
> summary(res.aov)
            Df Sum Sq Mean Sq F value   Pr(>F)
gtype        5  979.4  195.88   53.62 8.57e-08 ***
Residuals   12   43.8    3.65
> TukeyHSD(res.aov)
  Tukey multiple comparisons of means
    95% family-wise confidence level
Fit: aov(formula = medlife ~ gtype, data = df)

                          diff         lwr        upr     p adj
dnr1[2-133]-AMP14    10.833333   5.5917105  16.074956 0.0001753 ***
iso-AMP14            14.333333   9.0917105  19.574956 0.0000104 ***
mth-AMP14            13.000000   7.7583771  18.241623 0.0000286 ***
OR-R-AMP14           15.333333  10.0917105  20.574956 0.0000051 ***
Rel-AMP14            -3.666667  -8.9082895   1.574956 0.2473254 ns
iso-dnr1[2-133]       3.500000  -1.7416229   8.741623 0.2874264 ns
mth-dnr1[2-133]       2.166667  -3.0749562   7.408290 0.7330487 ns
OR-R-dnr1[2-133]      4.500000  -0.7416229   9.741623 0.1093032 ns
Rel-dnr1[2-133]     -14.500000 -19.7416229  -9.258377 0.0000092 ***
mth-iso              -1.333333  -6.5749562   3.908290 0.9505062 ns
OR-R-iso              1.000000  -4.2416229   6.241623 0.9852874 ns
Rel-iso             -18.000000 -23.2416229 -12.758377 0.0000009 ***
OR-R-mth              2.333333  -2.9082895   7.574956 0.6734614 ns
Rel-mth             -16.666667 -21.9082895 -11.425044 0.0000021 ***
Rel-OR-R            -19.000000 -24.2416229 -13.758377 0.0000005 ***
```

*dnr1 05-04-2022 median calc'd as female - avg(female-male difference)/2 of previous experiments = 38-3=35
