## Supplementary material for "Antimicrobial peptides do not directly contribute to aging in *Drosophila*, but improve lifespan by preventing dysbiosis": File S5: Cox_29C_mf.R

```
### Useful link http://anson.ucdavis.edu/~hiwang/teaching/10fall/R_tutorial%201.pdf

setwd("~/Desktop/Lemaitre Lab/Aging paper OFFLINE/Revisions/Figures_SuppFiles/FileS5_rawdata/FigS1")
library(survival) # R only
#install.packages("print.matrix")
library(colorspace)
#pal<-choose_palette()
library(hues)

library(readxl)
#FlyData <- read_excel("/Volumes/Lemaitre-Lab/Mark Hanson/Mark-Infections/2020/Longevity/Baramycin CoxPH/Baramycin-CoxPH_mf.xls")
FlyData <- read_excel("~/Desktop/Lemaitre Lab/Aging paper OFFLINE/Revisions/Figures_SuppFiles/FileS5_rawdata/FigS1/Cox_29C_mf.xls")
#FlyData <- read_excel("/Volumes/Lemaitre-Lab/Mark Hanson/Mark-Infections/2020/Longevity/Baramycin CoxPH/Baramycin-CoxPH_males.xlsx")
#FlyData <- read.csv(file.choose(), header=T)
head(FlyData)

#########################################################
###Adding to get median lifespans ***PER EXPERIMENT***###
#########################################################
#GtypeExp<-paste(FlyData$Genotype,FlyData$Experiment,FlyData$Sex)
#FlyData$Genotype<-GtypeExp

Gtype<-as.factor(FlyData$Genotype) #Note: it will organize levels alphabetically first by capitals then by lower case
levels(Gtype)
Gtype2<-unique(c(as.character(levels(Gtype)))) #Sorts genotypes by alphabetical order, doesn't require you to enter them in alphabetical order!
Life<-FlyData$Time2 #Life where time can't be -40000ish
Status<-FlyData$Censored #Censoring status, 0=censored
Sex<-FlyData$Sex
Exp<-FlyData$Experiment

###################################################
#Convert R survival data to Graphpad format
#Answer provided by Sotos in https://stackoverflow.com/questions/47770488/like-dcast-but-without-sum-of-data

###Prepare dataframe for export###
FlyData$Time.dead <- NULL #Get rid of extraneous column
Gtype<-as.factor(FlyData$Genotype); Life<-FlyData$Time2 ; Status<-as.numeric(FlyData$Censored)
FlyData<-data.frame(Gtype,Life,Status)

###Convert to Graphpad format###
#install.packages("backports")
library(backports)
library(tidyverse)
df<-FlyData %>% mutate(new = row_number()) %>% spread(Gtype, Status, fill = NA) %>% select(-new)
#write.csv(df, file="Baramycin-CoxPH_females.csv")
write.csv(df, file="Cox_29C_mf.csv")
#write.csv(df, file="Baramycin-CoxPH_mf.csv")
###################################################

### Kaplan-Meier test statistics and survival plot separated by Treatment
quartz(h=6,w=9) ## The equivalent for a PC is windows()
par(mfrow=c(1,1),mar=par()$mar+c(0,0,0,7), pty="s", xpd=TRUE)
flysurv <- survfit( Surv(Life, Status)~ strata(Gtype))
plot(flysurv, lty=c(1:3), lwd=c(1.5), col=iwanthue(20), xlab="Time (days)", ylab="Survival Probability", mark=c(1:20), xaxt="n", yaxt="n")#conf.times=c(1,2,3,4,5,6,7), conf.cap=.005
points(flysurv,col=iwanthue(20),pch=c(1:20))
axis(side=1,at=seq(0,50,by=5), cex.axis=0.8)
axis(side=2,at=seq(0,1,by=.25))
legend(55,1, (as.factor(Gtype2)), lty=c(1:3),lwd=c(1.5), col=iwanthue(20), cex = 1, pch=c(1:20)) #set legend to x=16 for graphs with xlim 14, and 8 for graphs with xlim 7
title(main="Longevity, 29'C, Males + Females
2022 three experiments",cex.main=0.8,line=1)
text(90,0,"coxph(formula = Surv(Life, Status) ~ Gtype + Exp + Sex, method = efron)
iso as baseline,  n= 707, number of events= 680

                         coef exp(coef)  se(coef)      z Pr(>|z|)    
Gtype2. OR-R         -0.27262   0.76138   0.13749 -1.983   0.0474 *  
Gtype3. AMP14         3.55857  35.11296   0.19711 18.053  < 2e-16 ***
Gtype4. iso Rel       5.60376 271.44474   0.25954 21.591  < 2e-16 ***
Gtype5. mth[1]        0.34363   1.41005   0.13249  2.594   0.0095 ** 
Gtype6. dnr1[2-133]   0.92560   2.52339   0.13997  6.613 3.77e-11 ***
Exp***, Sex**


Call: survfit(formula = Surv(Life, Status) ~ strata(Gtype))

                                         n events median 0.95LCL 0.95UCL
strata(Gtype)=1. iso 05-04-2022         40     36   38.0      36      41
strata(Gtype)=1. iso 20-03-2022         40     40   38.0      38      41
strata(Gtype)=1. iso 24-03-2022         40     40   38.0      34      43
strata(Gtype)=2. OR-R 05-04-2022        40     23   41.0      34      NA
strata(Gtype)=2. OR-R 20-03-2022        40     36   38.0      38      43
strata(Gtype)=2. OR-R 24-03-2022        38     34   41.0      34      43
strata(Gtype)=3. AMP14 05-04-2022       30     30   21.0      20      NA
strata(Gtype)=3. AMP14 20-03-2022       39     39   27.0      27      27
strata(Gtype)=3. AMP14 24-03-2022       40     40   23.0      23      25
strata(Gtype)=4. iso Rel 05-04-2022     40     40   20.0      NA      NA
strata(Gtype)=4. iso Rel 20-03-2022     40     40   22.0      NA      NA
strata(Gtype)=4. iso Rel 24-03-2022     40     40   18.0      NA      NA
strata(Gtype)=5. mth[1] 05-04-2022      40     39   38.0      36      NA
strata(Gtype)=5. mth[1] 20-03-2022      40     40   38.0      38      41
strata(Gtype)=5. mth[1] 24-03-2022      40     40   34.0      34      37
strata(Gtype)=6. dnr1[2-133] 05-04-2022 40     24   35.0*      36      NA
strata(Gtype)=6. dnr1[2-133] 20-03-2022 40     40   34.5      31      38
strata(Gtype)=6. dnr1[2-133] 24-03-2022 40     40   34.0      34      37

*dnr1 05-04-2022 median calc'd as female - avg(female-male difference)/2 of previous experiments = 38-3=35",cex=0.48,pos=2) #set x to 25 for graphs with xlim = c(0,14), and 12.5 for graphs with xlim = c(0,7)
text(-6,-.1,"> summary(res.aov)
            Df Sum Sq Mean Sq F value   Pr(>F)    
gtype        5  979.4  195.88   53.62 8.57e-08 ***
Residuals   12   43.8    3.65                     
> TukeyHSD(res.aov)
  Tukey multiple comparisons of means
    95% family-wise confidence level
Fit: aov(formula = medlife ~ gtype, data = df)

                        diff         lwr        upr     p adj
dnr1[2-133]-AMP14  10.833333   5.5917105  16.074956 0.0001753 ***
iso-AMP14          14.333333   9.0917105  19.574956 0.0000104 ***
mth-AMP14          13.000000   7.7583771  18.241623 0.0000286 ***
OR-R-AMP14         15.333333  10.0917105  20.574956 0.0000051 ***
Rel-AMP14          -3.666667  -8.9082895   1.574956 0.2473254 ns
iso-dnr1[2-133]     3.500000  -1.7416229   8.741623 0.2874264 ns
mth-dnr1[2-133]     2.166667  -3.0749562   7.408290 0.7330487 ns
OR-R-dnr1[2-133]    4.500000  -0.7416229   9.741623 0.1093032 ns
Rel-dnr1[2-133]   -14.500000 -19.7416229  -9.258377 0.0000092 ***
mth-iso            -1.333333  -6.5749562   3.908290 0.9505062 ns
OR-R-iso            1.000000  -4.2416229   6.241623 0.9852874 ns
Rel-iso           -18.000000 -23.2416229 -12.758377 0.0000009 ***
OR-R-mth            2.333333  -2.9082895   7.574956 0.6734614 ns
Rel-mth           -16.666667 -21.9082895 -11.425044 0.0000021 ***
Rel-OR-R          -19.000000 -24.2416229 -13.758377 0.0000005 ***",cex=0.28,pos=2) #set x to 25 for graphs with xlim = c(0,14), and 12.5 for graphs with xlim = c(0,7)

fix.coxph <- coxph(Surv(Life, Status)~Gtype+Exp,method='efron')
summary(fix.coxph)
#This says dnr1 highly different, but not OR-R (p = .076)

int.coxph <- coxph(Surv(Life, Status)~Gtype*Exp*Sex,method='efron')
summary(int.coxph)
#This says dnr1 not really different, but that OR-R is (p < .001, because OR-R has consistent sex effects across experiments while iso didn't. iso uniquely a bad reference genotype for the sex effect. Could possibly reveal this using a separate column -vial- that becomes redundant to the sex*exp interaction)

###########################################################################
#SEE BELOW#
###########################################################################
#Call: survfit(formula = Surv(Life, Status) ~ strata(Gtype))
#   13 observations deleted due to missingness 
#                                           n events median 0.95LCL 0.95UCL
#strata(Gtype)=1. iso 05-04-2022 f         20     19   36.0      34      38
#strata(Gtype)=1. iso 05-04-2022 m         20     17   41.0      38      41
#strata(Gtype)=1. iso 20-03-2022 f         20     20   38.0      38      43
#strata(Gtype)=1. iso 20-03-2022 m         20     20   38.0      38      41
#strata(Gtype)=1. iso 24-03-2022 f         20     20   43.0      41      NA
#strata(Gtype)=1. iso 24-03-2022 m         20     20   34.0      34      39
#strata(Gtype)=2. OR-R 05-04-2022 f        20      6     NA      NA      NA
#strata(Gtype)=2. OR-R 05-04-2022 m        20     17   34.0      30      36
#strata(Gtype)=2. OR-R 20-03-2022 f        20     18   43.0      41      45
#strata(Gtype)=2. OR-R 20-03-2022 m        20     18   38.0      33      41
#strata(Gtype)=2. OR-R 24-03-2022 f        18     18   43.0      41      46
#strata(Gtype)=2. OR-R 24-03-2022 m        20     16   34.0      29      37
#strata(Gtype)=3. AMP14 05-04-2022 f       14     14   25.0      22      NA
#strata(Gtype)=3. AMP14 05-04-2022 m       16     16   20.0      20      NA
#strata(Gtype)=3. AMP14 20-03-2022 f       19     19   27.0      27      29
#strata(Gtype)=3. AMP14 20-03-2022 m       20     20   27.0      NA      NA
#strata(Gtype)=3. AMP14 24-03-2022 f       20     20   26.0      23      27
#strata(Gtype)=3. AMP14 24-03-2022 m       20     20   23.0      23      23
#strata(Gtype)=4. iso Rel 05-04-2022 f     20     20   20.0      NA      NA
#strata(Gtype)=4. iso Rel 05-04-2022 m     20     20   20.0      NA      NA
#strata(Gtype)=4. iso Rel 20-03-2022 f     20     20   22.0      NA      NA
#strata(Gtype)=4. iso Rel 20-03-2022 m     20     20   22.0      19      NA
#strata(Gtype)=4. iso Rel 24-03-2022 f     20     20   18.0      NA      NA
#strata(Gtype)=4. iso Rel 24-03-2022 m     20     20   18.0      NA      NA
#strata(Gtype)=5. mth[1] 05-04-2022 f      20     20   39.5      36      NA
#strata(Gtype)=5. mth[1] 05-04-2022 m      20     19   38.0      36      NA
#strata(Gtype)=5. mth[1] 20-03-2022 f      20     20   43.0      38      43
#strata(Gtype)=5. mth[1] 20-03-2022 m      20     20   38.0      36      38
#strata(Gtype)=5. mth[1] 24-03-2022 f      20     20   37.0      34      41
#strata(Gtype)=5. mth[1] 24-03-2022 m      20     20   34.0      34      37
#strata(Gtype)=6. dnr1[2-133] 05-04-2022 f 20     18   38.0      36      NA
#strata(Gtype)=6. dnr1[2-133] 05-04-2022 m 20      6     NA      NA      NA
#strata(Gtype)=6. dnr1[2-133] 20-03-2022 f 20     20   38.0      38      38
#strata(Gtype)=6. dnr1[2-133] 20-03-2022 m 20     20   29.5      27      33
#strata(Gtype)=6. dnr1[2-133] 24-03-2022 f 20     20   35.5      34      NA
#strata(Gtype)=6. dnr1[2-133] 24-03-2022 m 20     20   32.0      32      34
############################################################################
#dnr1-m_05-04-2022 censored 25dpi thus NA
############################################################################

flysurv

survoutput<-summary(flysurv)$table
write.csv(survoutput,file="survoutput.csv")


##
#Testing contribution of frailty()
##
###https://app.datacamp.com/workspace/preview?_tag=rdocs&rdocsPath=packages%2Fsurvival%2Fversions%2F2.26%2Ftopics%2Ffrailty&utm_source=r-docs&utm_medium=docs&utm_term=frailty&utm_content=run_example_in_workspace
###FRAILTY function considered for random effect
time<-Life
status<-Status
kfit <- coxph(Surv(time, status)~ Gtype + Sex + frailty(Exp), FlyData)
summary(kfit)
kfit0 <- coxph(Surv(time, status)~ Gtype + Sex + frailty(Exp), FlyData)
summary(kfit0)
kfitm1 <- coxph(Surv(time,status)~ Gtype + Sex + frailty(Exp, dist='gauss'), FlyData)
summary(kfitm1)
CoxAIC<-coxph(Surv(time,status)~ Gtype + Sex + frailty(Exp, dist='gauss', method='aic',caic=TRUE), FlyData)
summary(CoxAIC)
# uncorrected aic
CoxAIC2<-coxph(Surv(time,status)~ Gtype + Sex + frailty(Exp, method='aic', caic=FALSE), FlyData)
summary(CoxAIC2)

###
#All of thesefrailty examples illustrate how unnecessary it is to include in the model
###


#############################################
#Some t.tests for CR vs ABX median lifespans#
#############################################

iso<-c(38,38,38)
ORR<-c(38,41,41)
AMP14<-c(21,27,23)
Rel<-c(20,22,18)
mth<-c(38,34,38)
dnr1<-c(35,34.5,34) #value from 5/4/22 entered as 35, needs to be adjusted from observations because of early male censorship, f avg 6 days later than m avg in prev. exp... so avg 3 above median proper

gtype<-c("iso","iso","iso","OR-R","OR-R","OR-R","AMP14","AMP14","AMP14","Rel","Rel","Rel","mth","mth","mth","dnr1[2-133]","dnr1[2-133]","dnr1[2-133]")
medlife<-c(38,38,38,38,41,38,21,27,23,20,22,18,38,38,34,35,34.5,34)

df<-data.frame(gtype,medlife)

# Compute the analysis of variance
res.aov <- aov(medlife ~ gtype,data=df)
# Summary of the analysis
summary(res.aov)
TukeyHSD(res.aov)

###########################
#PRELIMINARY one-way ANOVA using estimated iso and OR-R values for remaining experiments
###########################
#> TukeyHSD(res.aov)
#  Tukey multiple comparisons of means
#    95% family-wise confidence level
#
#Fit: aov(formula = medlife ~ gtype, data = df)
#
#$gtype
#                        diff         lwr        upr     p adj sig
#dnr1[2-133]-AMP14   7.500000  -0.4302365  15.430237 0.0677353 .
#iso-AMP14          13.833333   5.9030968  21.763570 0.0008426 ***
#mth-AMP14          12.333333   4.4030968  20.263570 0.0022602 **
#OR-R-AMP14         19.333333  11.4030968  27.263570 0.0000341 ***
#Rel-AMP14          -3.666667 -11.5969032   4.263570 0.6404264 
#iso-dnr1[2-133]     6.333333  -1.5969032  14.263570 0.1502391 
#mth-dnr1[2-133]     4.833333  -3.0969032  12.763570 0.3727122 
#OR-R-dnr1[2-133]   11.833333   3.9030968  19.763570 0.0031725 **
#Rel-dnr1[2-133]   -11.166667 -19.0969032  -3.236430 0.0050219 **
#mth-iso            -1.500000  -9.4302365   6.430237 0.9858339 
#OR-R-iso            5.500000  -2.4302365  13.430237 0.2545197 
#Rel-iso           -17.500000 -25.4302365  -9.569763 0.0000926 ***
#OR-R-mth            7.000000  -0.9302365  14.930237 0.0958691 .
#Rel-mth           -16.000000 -23.9302365  -8.069763 0.0002207 ***
#Rel-OR-R          -23.000000 -30.9302365 -15.069763 0.0000056 ***


#####################################
#Multiple comparisons via Robyn code#
#####################################

m<-c()
for (i in 1:length(levels(Gtype))){
  levels<-c(levels(Gtype)[i], levels(Gtype)[-i])
  df<-data.frame(Life=Life, Status=Status, Gtype_prime=Gtype)
  df$Gtype_prime<-factor(df$Gtype_prime, levels=levels)
  full.coxph <- coxph(Surv(df$Life, df$Status)~df$Gtype_prime,method='exact')
  s<-summary(full.coxph)
  p<-as.numeric(s$coefficients[,5])
  p2<-c(p, NA)
  if (i!=length(levels(Gtype))){
    for (j in rev((i+1):length(levels(Gtype)))){
      p2[j]<-p2[j-1]
      
    }
  }
  p2[i]<-NA
  m<-c(m, p2)
}

#quartz(h=6,w=9) ## The equivalent for a PC is windows()
#par(mfrow=c(1,1),mar=par()$mar+c(0,0,0,7), pty="s", xpd=TRUE)
Results<-matrix(m, nrow=length(levels(Gtype)))
colnames(Results)<-levels(Gtype)
rownames(Results)<-levels(Gtype)
library(plot.matrix)
Results_v2<-Results
Results_v2[Results_v2>0.05]<-NA
#plot(Results_v2, cex.axis=0.4,las = 2, ylab="", xlab="")

write.csv(Results_v2, file="Results_v2.csv")
```
