## Supplementary material for "Antimicrobial peptides do not directly contribute to aging in *Drosophila*, but improve lifespan by preventing dysbiosis": File S5: Act-Repo-Gal4_CoxPH.pdf

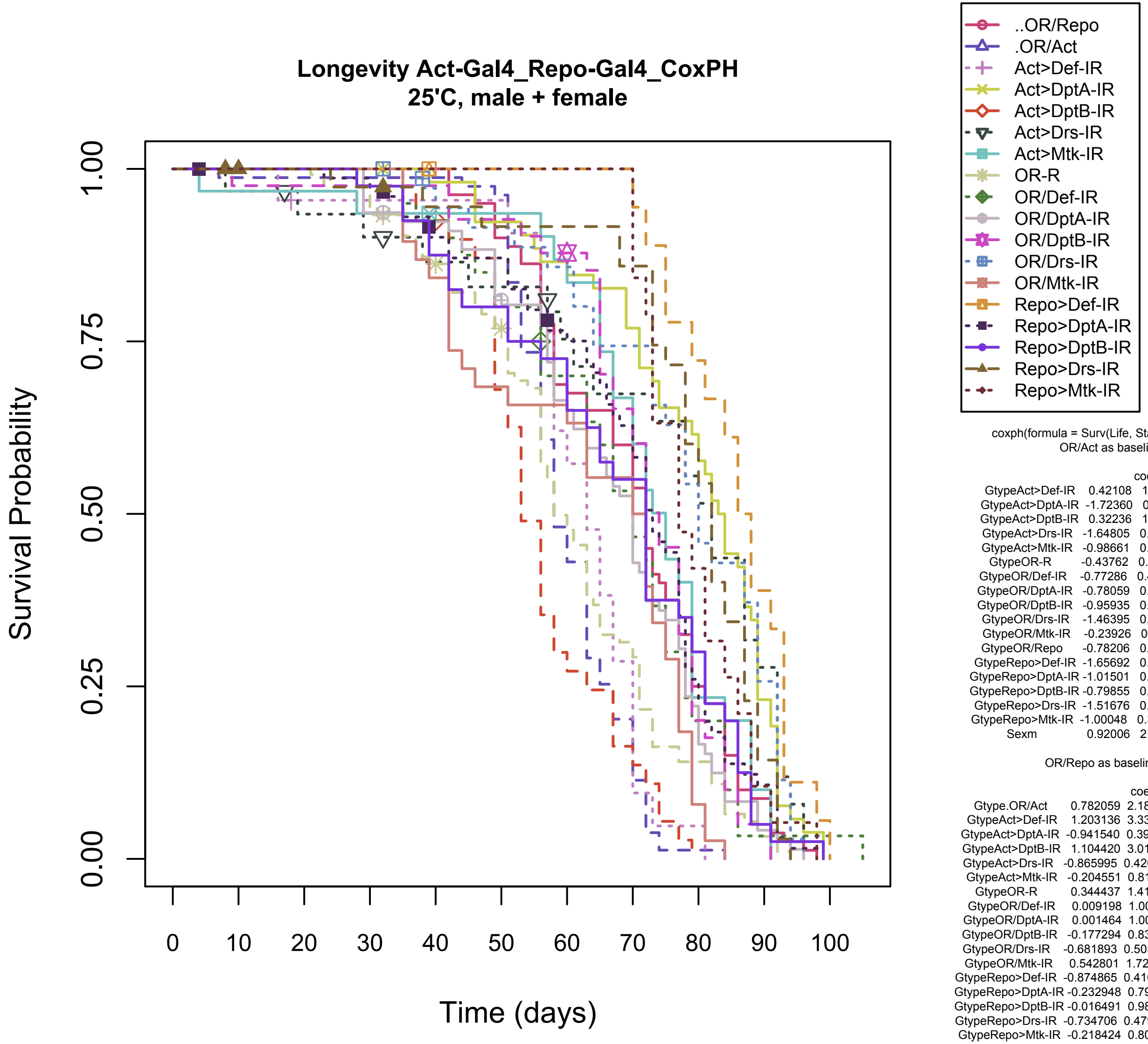


```
coxph(formula = Surv(Life, Status) ~ Gtype+Sex, method = efron)
OR/Act as baseline,  n= 890, number of events= 817

                       coef exp(coef) se(coef)      z Pr(>|z|)
GtypeAct>Def-IR     0.42108   1.52360  0.25316  1.663 0.096255 .
GtypeAct>DptA-IR   -1.72360   0.17842  0.18742 -9.196  < 2e-16 ***
GtypeAct>DptB-IR    0.32236   1.38038  0.20029  1.609 0.107512
GtypeAct>Drs-IR    -1.64805   0.19242  0.23258 -7.086 1.38e-12 ***
GtypeAct>Mtk-IR    -0.98661   0.37284  0.21770 -4.532 5.84e-06 ***
GtypeOR-R          -0.43762   0.64557  0.15467 -2.829 0.004665 **
GtypeOR/Def-IR     -0.77286   0.46169  0.21554 -3.586 0.000336 ***
GtypeOR/DptA-IR    -0.78059   0.45813  0.16644 -4.690 2.73e-06 ***
GtypeOR/DptB-IR    -0.95935   0.38314  0.19701 -4.870 1.12e-06 ***
GtypeOR/Drs-IR     -1.46395   0.23132  0.21041 -6.958 3.46e-12 ***
GtypeOR/Mtk-IR     -0.23926   0.78721  0.20111 -1.190 0.234162
GtypeOR/Repo       -0.78206   0.45746  0.16476 -4.747 2.07e-06 ***
GtypeRepo>Def-IR   -1.65692   0.19072  0.27313 -6.066 1.31e-09 ***
GtypeRepo>DptA-IR  -1.01501   0.36240  0.17214 -5.897 3.71e-09 ***
GtypeRepo>DptB-IR  -0.79855   0.44998  0.19979 -3.997 6.41e-05 ***
GtypeRepo>Drs-IR   -1.51676   0.21942  0.20843 -7.277 3.41e-13 ***
GtypeRepo>Mtk-IR   -1.00048   0.36770  0.26437 -3.784 0.000154 ***
Sexm                0.92006   2.50945  0.07967 11.549  < 2e-16 ***

OR/Repo as baseline,  n= 890, number of events= 817

                        coef exp(coef)  se(coef)      z Pr(>|z|)
Gtype.OR/Act        0.782059  2.185968  0.164763  4.747 2.07e-06 ***
GtypeAct>Def-IR     1.203136  3.330544  0.249577  4.821 1.43e-06 ***
GtypeAct>DptA-IR   -0.941540  0.390027  0.182725 -5.153 2.57e-07 ***
GtypeAct>DptB-IR    1.104420  3.017475  0.202817  5.445 5.17e-08 ***
GtypeAct>Drs-IR    -0.865995  0.420633  0.228624 -3.788 0.000152 ***
GtypeAct>Mtk-IR    -0.204551  0.815013  0.215022 -0.951 0.341452
GtypeOR-R           0.344437  1.411195  0.153718  2.241 0.025045 *
GtypeOR/Def-IR      0.009198  1.009240  0.211844  0.043 0.965369
GtypeOR/DptA-IR     0.001464  1.001465  0.163141  0.009 0.992841
GtypeOR/DptB-IR    -0.177294  0.837534  0.195829 -0.905 0.365280
GtypeOR/Drs-IR     -0.681893  0.505659  0.203829 -3.345 0.000822 ***
GtypeOR/Mtk-IR      0.542801  1.720819  0.198504  2.734 0.006248 **
GtypeRepo>Def-IR   -0.874865  0.416918  0.264432 -3.308 0.000938 ***
GtypeRepo>DptA-IR  -0.232948  0.792195  0.167461 -1.391 0.164209
GtypeRepo>DptB-IR  -0.016491  0.983644  0.194146 -0.085 0.932308
GtypeRepo>Drs-IR   -0.734706  0.479646  0.204767 -3.588 0.000333 ***
GtypeRepo>Mtk-IR   -0.218424  0.803784  0.256443 -0.852 0.394355
Sexm                0.920065  2.509453  0.079669 11.549  < 2e-16 ***
```
