## Supplementary material for "Antimicrobial peptides do not directly contribute to aging in *Drosophila*, but improve lifespan by preventing dysbiosis": File S5: Repo mf CoxPH.pdf

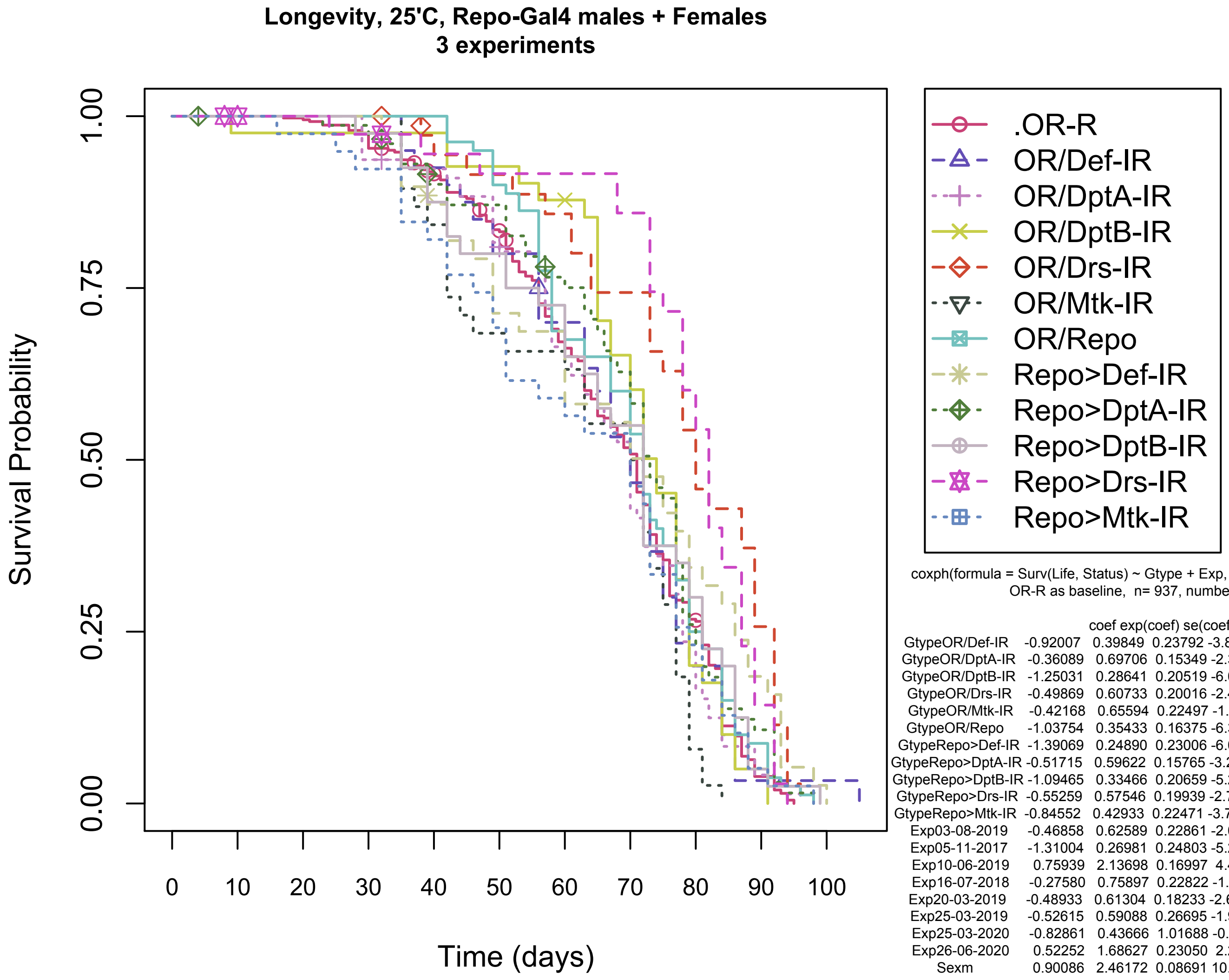


coxph(formula = Surv(Life, Status) ~ Gtype + Exp, method = efron)
OR-R as baseline,  n= 937, number of events= 814

| | coef | exp(coef) | se(coef) | z | Pr(>\|z\|) | |
|---|---|---|---|---|---|---|
| GtypeOR/Def-IR | -0.92007 | 0.39849 | 0.23792 | -3.867 | 0.000110 | *** |
| GtypeOR/DptA-IR | -0.36089 | 0.69706 | 0.15349 | -2.351 | 0.018716 | * |
| GtypeOR/DptB-IR | -1.25031 | 0.28641 | 0.20519 | -6.093 | 1.11e-09 | *** |
| GtypeOR/Drs-IR | -0.49869 | 0.60733 | 0.20016 | -2.491 | 0.012723 | * |
| GtypeOR/Mtk-IR | -0.42168 | 0.65594 | 0.22497 | -1.874 | 0.060872 | . |
| GtypeOR/Repo | -1.03754 | 0.35433 | 0.16375 | -6.336 | 2.36e-10 | *** |
| GtypeRepo>Def-IR | -1.39069 | 0.24890 | 0.23006 | -6.045 | 1.50e-09 | *** |
| GtypeRepo>DptA-IR | -0.51715 | 0.59622 | 0.15765 | -3.280 | 0.001037 | ** |
| GtypeRepo>DptB-IR | -1.09465 | 0.33466 | 0.20659 | -5.299 | 1.17e-07 | *** |
| GtypeRepo>Drs-IR | -0.55259 | 0.57546 | 0.19939 | -2.771 | 0.005581 | ** |
| GtypeRepo>Mtk-IR | -0.84552 | 0.42933 | 0.22471 | -3.763 | 0.000168 | *** |
| Exp03-08-2019 | -0.46858 | 0.62589 | 0.22861 | -2.050 | 0.040395 | * |
| Exp05-11-2017 | -1.31004 | 0.26981 | 0.24803 | -5.282 | 1.28e-07 | *** |
| Exp10-06-2019 | 0.75939 | 2.13698 | 0.16997 | 4.468 | 7.90e-06 | *** |
| Exp16-07-2018 | -0.27580 | 0.75897 | 0.22822 | -1.208 | 0.226875 | |
| Exp20-03-2019 | -0.48933 | 0.61304 | 0.18233 | -2.684 | 0.007280 | ** |
| Exp25-03-2019 | -0.52615 | 0.59088 | 0.26695 | -1.971 | 0.048727 | * |
| Exp25-03-2020 | -0.82861 | 0.43666 | 1.01688 | -0.815 | 0.415153 | |
| Exp26-06-2020 | 0.52252 | 1.68627 | 0.23050 | 2.267 | 0.023396 | * |
| Sexm | 0.90086 | 2.46172 | 0.08691 | 10.366 | < 2e-16 | *** |
