## Supplementary material for "Antimicrobial peptides do not directly contribute to aging in *Drosophila*, but improve lifespan by preventing dysbiosis": File S5: Repo_Rel-IR_mf.pdf

### Longevity, Repo>Rel-IR, Males + Females
## 20-03-2022, 25-03-2022, 08-04-2022
### 2019 experiment removed because did not share wt control for crosses, but same direction seen

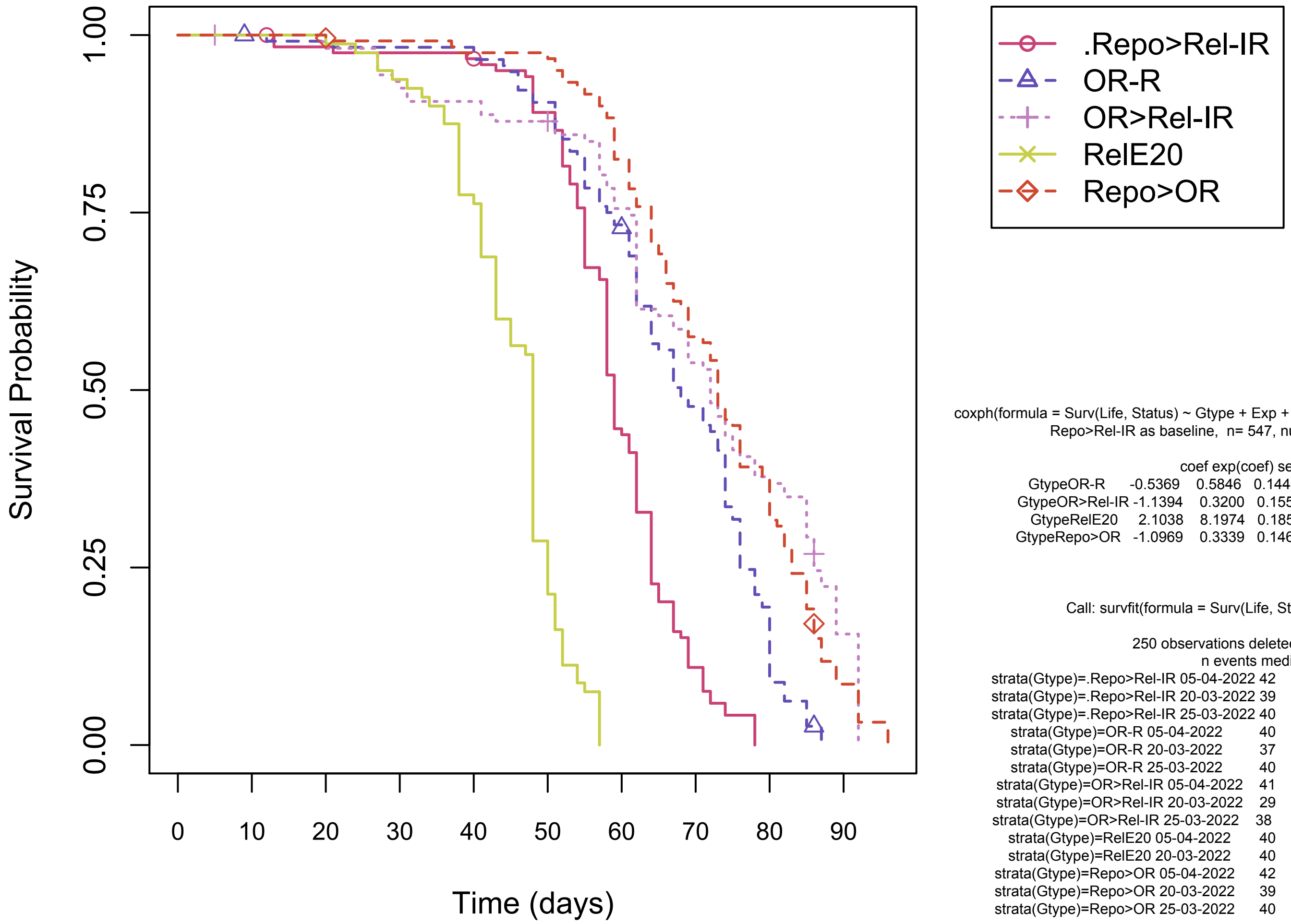


```
coxph(formula = Surv(Life, Status) ~ Gtype + Exp + Sex, method = efron)
Repo>Rel-IR as baseline,  n= 547, number of events= 518

                     coef exp(coef) se(coef)      z Pr(>|z|)
GtypeOR-R         -0.5369    0.5846   0.1444 -3.717 0.000201 ***
GtypeOR>Rel-IR    -1.1394    0.3200   0.1550 -7.351 1.97e-13 ***
GtypeRelE20        2.1038    8.1974   0.1853 11.353  < 2e-16 ***
GtypeRepo>OR      -1.0969    0.3339   0.1461 -7.506 6.10e-14 ***
                                                          Exp ***
                                                          Sex ***

Call: survfit(formula = Surv(Life, Status) ~ strata(Gtype))

250 observations deleted due to missingness
                                        n events median 0.95LCL 0.95UCL
strata(Gtype)=.Repo>Rel-IR 05-04-2022 42     42     59      55      62
strata(Gtype)=.Repo>Rel-IR 20-03-2022 39     37     64      59      67
strata(Gtype)=.Repo>Rel-IR 25-03-2022 40     40     58      58      61
strata(Gtype)=OR-R 05-04-2022         40     38     64      62      76
strata(Gtype)=OR-R 20-03-2022         37     37     68      64      78
strata(Gtype)=OR-R 25-03-2022         40     37     69      62      74
strata(Gtype)=OR>Rel-IR 05-04-2022    41     40     59      57      62
strata(Gtype)=OR>Rel-IR 20-03-2022    29     28     85      82      89
strata(Gtype)=OR>Rel-IR 25-03-2022    38     23     86      72      NA
strata(Gtype)=RelE20 05-04-2022       40     40     48      48      48
strata(Gtype)=RelE20 20-03-2022       40     40     43      43      50
strata(Gtype)=Repo>OR 05-04-2022      42     41     69      66      76
strata(Gtype)=Repo>OR 20-03-2022      39     39     80      73      87
strata(Gtype)=Repo>OR 25-03-2022      40     36     74      69      83
```
