## Supplementary material for "Antimicrobial peptides do not directly contribute to aging in *Drosophila*, but improve lifespan by preventing dysbiosis": File S5: Repo_Rel-IR_mf_iso.pdf

**Longevity, Repo>Rel-IR_iso, Males + Females**
**10-07-2019**

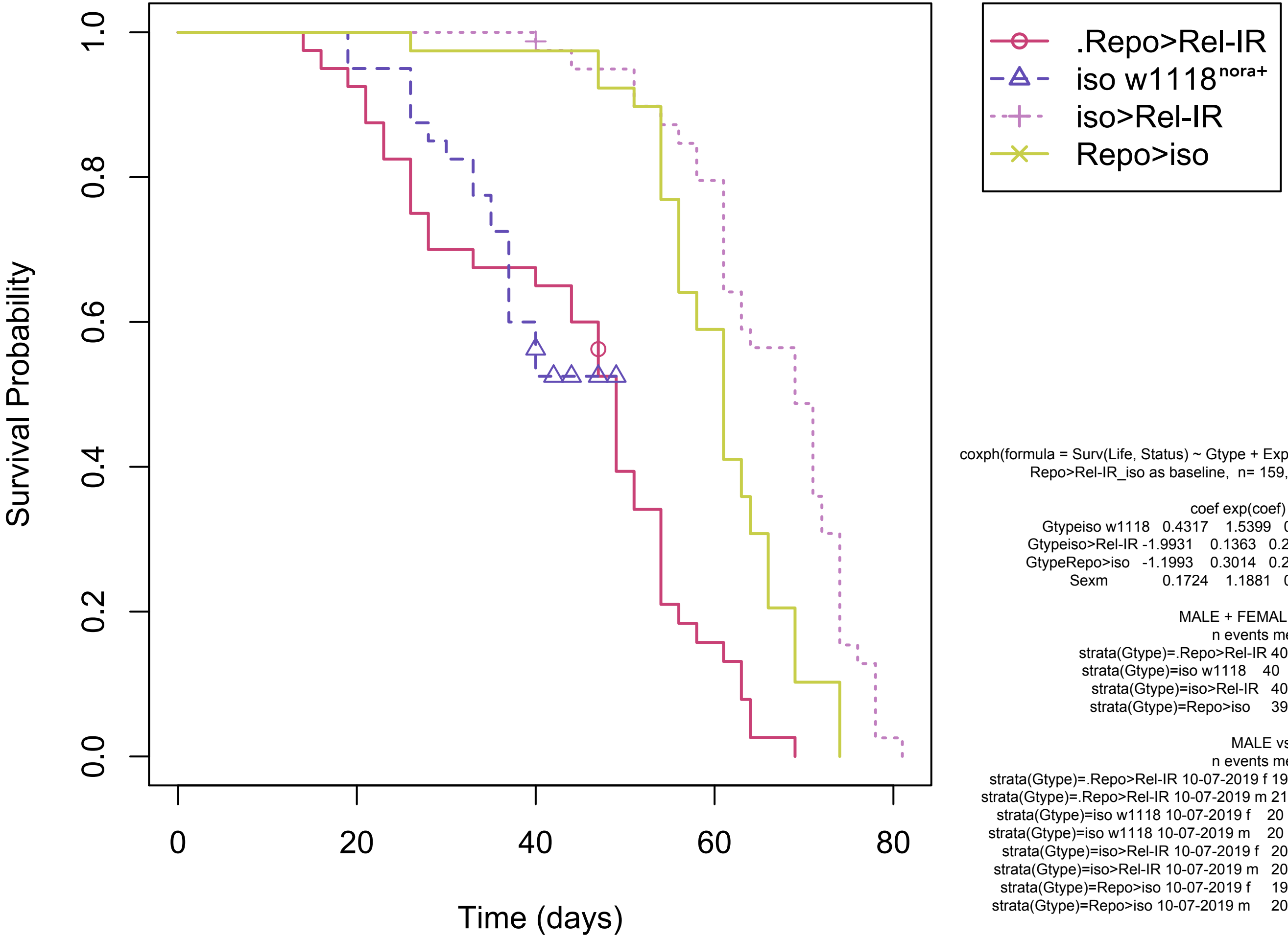


coxph(formula = Surv(Life, Status) ~ Gtype + Exp + Sex, method = efron)
Repo>Rel-IR_iso as baseline, n= 159, number of events= 136

| | coef | exp(coef) | se(coef) | z | Pr(>\|z\|) | |
|---|---|---|---|---|---|---|
| Gtypeiso w1118 | 0.4317 | 1.5399 | 0.3340 | 1.293 | 0.196 | |
| Gtypeiso>Rel-IR | -1.9931 | 0.1363 | 0.2726 | -7.311 | 2.66e-13 | *** |
| GtypeRepo>iso | -1.1993 | 0.3014 | 0.2384 | -5.031 | 4.87e-07 | *** |
| Sexm | 0.1724 | 1.1881 | 0.1784 | 0.966 | 0.334 | |

MALE + FEMALE MEDIAN LIFESPANS

| | n | events | median | 0.95LCL | 0.95UCL |
|---|---|---|---|---|---|
| strata(Gtype)=.Repo>Rel-IR | 40 | 39 | 49 | 44 | 54 |
| strata(Gtype)=iso w1118 | 40 | 19 | NA | 37 | NA |
| strata(Gtype)=iso>Rel-IR | 40 | 39 | 69 | 63 | 72 |
| strata(Gtype)=Repo>iso | 39 | 39 | 61 | 58 | 64 |

MALE vs. FEMALE LIFESPANS

| | n | events | median | 0.95LCL | 0.95UCL |
|---|---|---|---|---|---|
| strata(Gtype)=.Repo>Rel-IR 10-07-2019 f | 19 | 18 | 51 | 49 | 56 |
| strata(Gtype)=.Repo>Rel-IR 10-07-2019 m | 21 | 21 | 28 | 23 | 54 |
| strata(Gtype)=iso w1118 10-07-2019 f | 20 | 12 | 37 | 35 | NA |
| strata(Gtype)=iso w1118 10-07-2019 m | 20 | 7 | NA | 40 | NA |
| strata(Gtype)=iso>Rel-IR 10-07-2019 f | 20 | 19 | 72 | 69 | 78 |
| strata(Gtype)=iso>Rel-IR 10-07-2019 m | 20 | 20 | 62 | 61 | 71 |
| strata(Gtype)=Repo>iso 10-07-2019 f | 19 | 19 | 61 | 58 | 66 |
| strata(Gtype)=Repo>iso 10-07-2019 m | 20 | 20 | 61 | 56 | 69 |
