## Supplementary figures and images for "Antimicrobial peptides do not directly contribute to aging in *Drosophila*, but improve lifespan by preventing dysbiosis"

### Fig1A.pdf

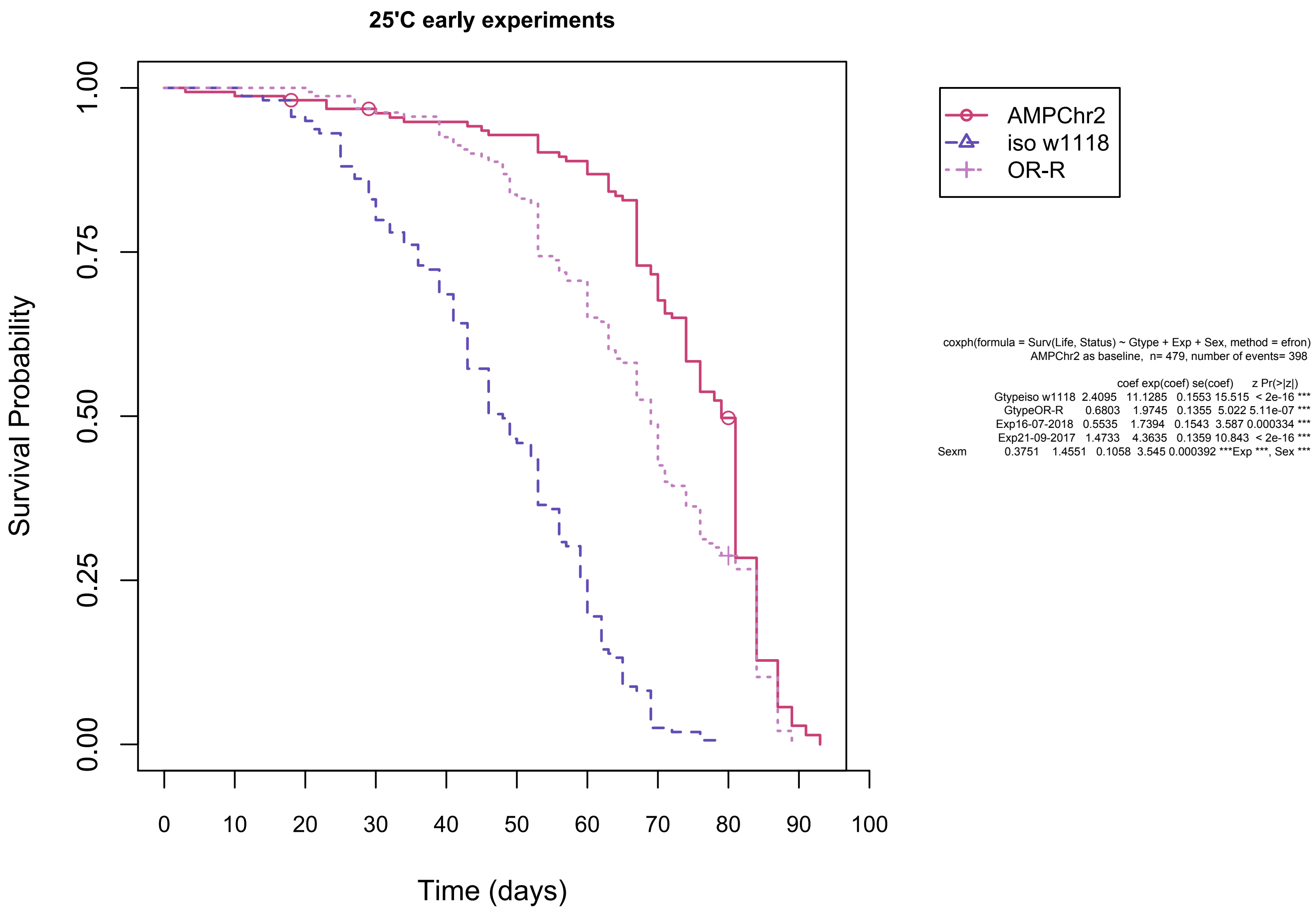

### iso nora example.pdf

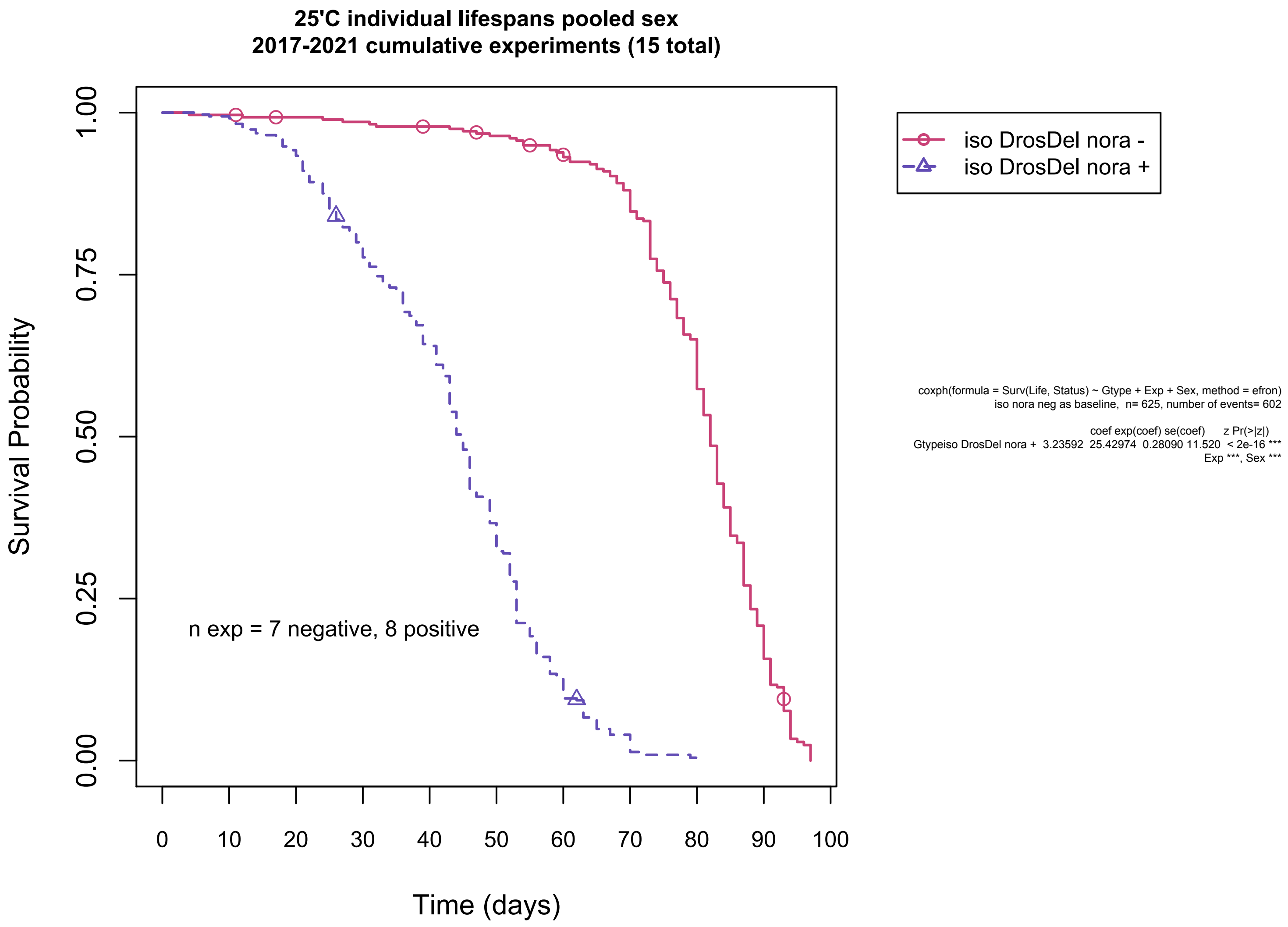

### Ligoxygakis exps.jpg

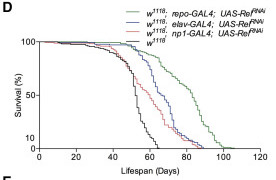
